# Glycolysis-stratified coordination of fatty acid and glutamine metabolism in pancreatic ductal adenocarcinoma

**DOI:** 10.1101/2025.09.14.676164

**Authors:** Salma Fareed

## Abstract

**Background:** Pancreatic ductal adenocarcinoma (PDAC) is highly lethal and characterized by profound metabolic rewiring. Prior work, including evidence that ketogenic diet (KD) exposure can increase tricarboxylic acid (TCA) activity and glutamine (GLN) dependence and sensitize PDAC to GLN-targeting strategies, suggests that fatty acid (FA) availability may modulate GLN utilization. However, it remains unclear how FA-GLN metabolic coordination varies across tumor glycolytic states and FA classes.

**Objective:** To quantify how FA-GLN coordination varies across PDAC glycolysis tertiles and FA classes (lipid families and chain-length enzyme classes).

**Methods:** RNA-seq and clinical data from a multi-institutional PDAC cohort (n=172) were analyzed to quantify FA and GLN pathway activities using single-sample gene set scoring (GSVA-derived pathway activity scores), followed by stratification of tumors into low, medium, and high glycolysis tertiles using ssGSEA-derived glycolysis scores. FA-GLN associations were evaluated at pathway, enzyme-class (medium-, long-, and very-long-chain FA; VLCFA ≥ C20), and family levels using correlation, Fisher-z meta-analysis, and bootstrap resampling. Family-level contrasts were used to compare FA-GLN coupling between glycolysis strata, and complementary metabolite-inference analyses were performed (n=173) to define metabolite-based tumor phenotypes. Protein-level validation was conducted in an independent PDAC proteomic cohort.

**Results:** FA-GLN coordination was strongly context-dependent: membrane-focused lipid families—glycerophospholipids (GPL) and sphingolipids (SP)—showed the strongest GLN coupling in glycolysis-low tumors, whereas coupling shifted toward VLCFA enzyme classes in glycolysis-medium tumors and weakened in glycolysis-high tumors. Family-level contrasts confirmed significantly weaker GPL–GLN and SP–GLN associations in high versus low glycolysis. Metabolite-inference analyses identified three metabolic phenotypes that preserved this glycolysis-stratified FA-GLN hierarchy, and protein-level analysis in the CPTAC PDAC cohort partially recapitulated preferential SP–GLN coordination in less glycolytic tumors. Survival analyses suggested glycolysis-state– and chain length-specific trends in FA-linked pathways.

**Conclusions:** Together, these findings delineate a glycolysis-stratified map of FA-GLN coordination in PDAC and nominate conditional metabolic vulnerabilities with potential therapeutic and dietary relevance.

Conceptual overview illustrating how coordinated fatty-acid and glutamine metabolism shifts across glycolysis-low, medium, and high PDAC tumors, highlighting FA family and chain length-specific metabolic dependencies.

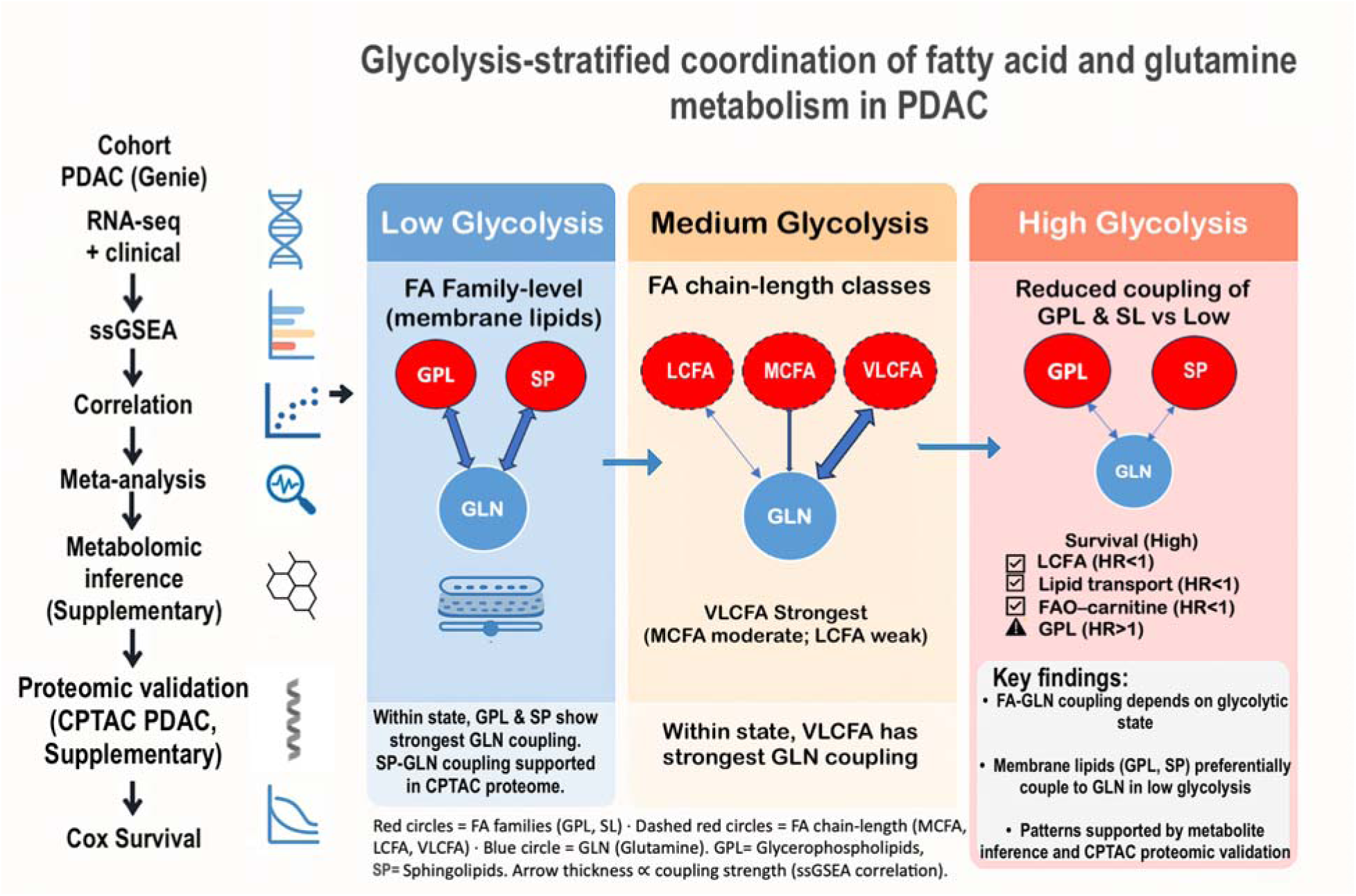

## Introduction

Pancreatic ductal adenocarcinoma (PDAC) is the most common form of pancreatic cancer and remains among the most lethal solid tumors. Despite progress in detection and therapy, the five-year survival rate remains below 12%, and incidence is projected to rise in the coming decades [1,2]. Most patients present with advanced or metastatic disease, and the limited benefit of current chemotherapies underscores the need for alternative therapeutic strategies [2]. In addition to genetic alterations, PDAC progression is shaped by metabolic adaptations that enable tumor cells to persist and proliferate in a hypoxic, nutrient-poor microenvironment [3,4]. Altered metabolism is recognized as a defining hallmark of malignancy, and PDAC is a typical example of metabolic plasticity [3,5]. The PDAC microenvironment is nutrient-limited, driving changes in nutrient use and energy pathways and fostering mechanisms such as extracellular protein scavenging [6]. In PDAC, glycolysis remains high despite oxygen availability, generating biosynthetic building blocks and redirecting metabolic dependencies [7]. Molecular subtyping indicates that glycolysis-high tumors tend to be more aggressive and linked to poorer clinical outcomes [8].

Glutamine (GLN) metabolism is central to PDAC biology. GLN provides carbon and nitrogen for nucleotide synthesis, sustains redox balance via glutathione, and replenishes the tricarboxylic acid (TCA) cycle through anaplerosis [9]. Oncogenic KRAS reprograms GLN utilization through non-canonical routes that support tumor growth [10]. Inhibition of GLN metabolism slows PDAC proliferation, but compensatory nutrient routes are frequently engaged [9,11]. Pharmacologic blockade with 6-diazo-5-oxo-L-norleucine (DON) has been shown to trigger a metabolic crisis in PDAC models, yet rapid adaptive responses were observed [12]. Stromal interactions also provide alternative substrates (e.g., alanine), complicating therapeutic targeting [13]. Direct metabolomic measurements on resected human PDAC tissue have demonstrated reduced glutamine relative to paired benign pancreas, underscoring nutrient limitation in the tumor microenvironment [6], emphasizing the importance of nutrient context.

Fatty acid (FA) metabolism likewise contributes substantially to PDAC. De novo lipogenesis and FA uptake furnish membrane components, signaling lipids, and energy reserves [14]. Enzymes that elongate and desaturate fatty acids (e.g., ELOVL, FADS, SCD) enable lipid remodeling and can promote oncogenic signaling [15]. PDAC cells also draw on exogenous lipids and alternative carbon sources such as acetate, reinforcing metabolic flexibility [16]. Distinct FA subclasses: medium-chain (MCFA), long-chain (LCFA), and very-long-chain fatty acids (VLCFA), likely exert different effects on tumor behavior, yet systematic, patient-based evaluation has been limited [17].

The interplay between FA and GLN metabolism in PDAC has been highlighted experimentally. Inhibition of GLN metabolism sensitizes tumors to ketogenic-diet conditions, revealing vulnerabilities when glucose and lipid pathways are simultaneously perturbed [18]. Additional work supports coordination between FA programs and GLN pathways and motivates combined therapeutic strategies targeting both axes [15,19]. However, FA metabolism is often treated as a single pathway, with limited attention to chain-length or family-specific diversity. Moreover, it remains unclear whether FA-GLN coordination is a general feature of PDAC or conditional on glycolytic activity. This gap is particularly relevant because ketogenic diets alter circulating FA chain-length distributions and patient tumors vary widely in glycolytic state [18].

To address these questions, transcriptomic and clinical data from a multi-institutional PDAC cohort (n=172) were analyzed. FA-GLN associations were evaluated across glycolytic strata, and chain-length-specific FA families were examined for prognostic and metabolic differences. A two-tier analytic framework was applied, combining broad profiling of curated FA pathways with targeted interrogation of MCFA, LCFA, and VLCFA families. Enzyme-class (chain-length) and family-level (functional) analyses were used to capture functional diversity, and associations with GLN were quantified using Elastic Net regression with bootstrap resampling. This study represents, to our knowledge, the first patient-based study to link FA-GLN coupling explicitly to glycolytic stratification in PDAC, delineating context-specific vulnerabilities with potential therapeutic and dietary relevance.

## Results

### FA-GLN correlations stratified by glycolysis state

Spearman correlations between fatty acid (FA) pathway activity and glutamine (GLN) activity differed by glycolysis state (Low, Medium, High glycolysis tertiles; ≈ 57-58 tumors per group; Fig 1, Table 1). In glycolysis-low tumors, sphingolipid pathways showed the strongest positive correlations with GLN, including Sphingolipid pathway (ρ = 0.44, FDR = 0.063), Sphingolipid metabolism overview (ρ = 0.39, FDR = 0.104), Sphingolipid de novo biosynthesis (ρ = 0.35, FDR = 0.104), and Sphingolipid metabolism integrated pathway (ρ = 0.35, FDR = 0.104). Glycerophospholipid biosynthesis also correlated positively (ρ = 0.35, FDR = 0.104). In glycolysis-medium tumors, the strongest correlations were observed for aminoacyl-tRNA pathways; tRNA aminoacylation (ρ = 0.46, FDR = 0.024), Mitochondrial tRNA aminoacylation (ρ = 0.45, FDR = 0.024), and Aminoacyl-tRNA biosynthesis (ρ = 0.42, FDR = 0.033). Lipid-associated pathways, including Peroxisomal lipid metabolism (ρ = 0.38, FDR = 0.078) and Mitochondrial fatty acid oxidation disorders (ρ = 0.36, FDR = 0.078), were also positively correlated. In glycolysis-high tumors, no FA pathways met the prespecified FDR ≤ 0.20 threshold.

**Fig 1.**
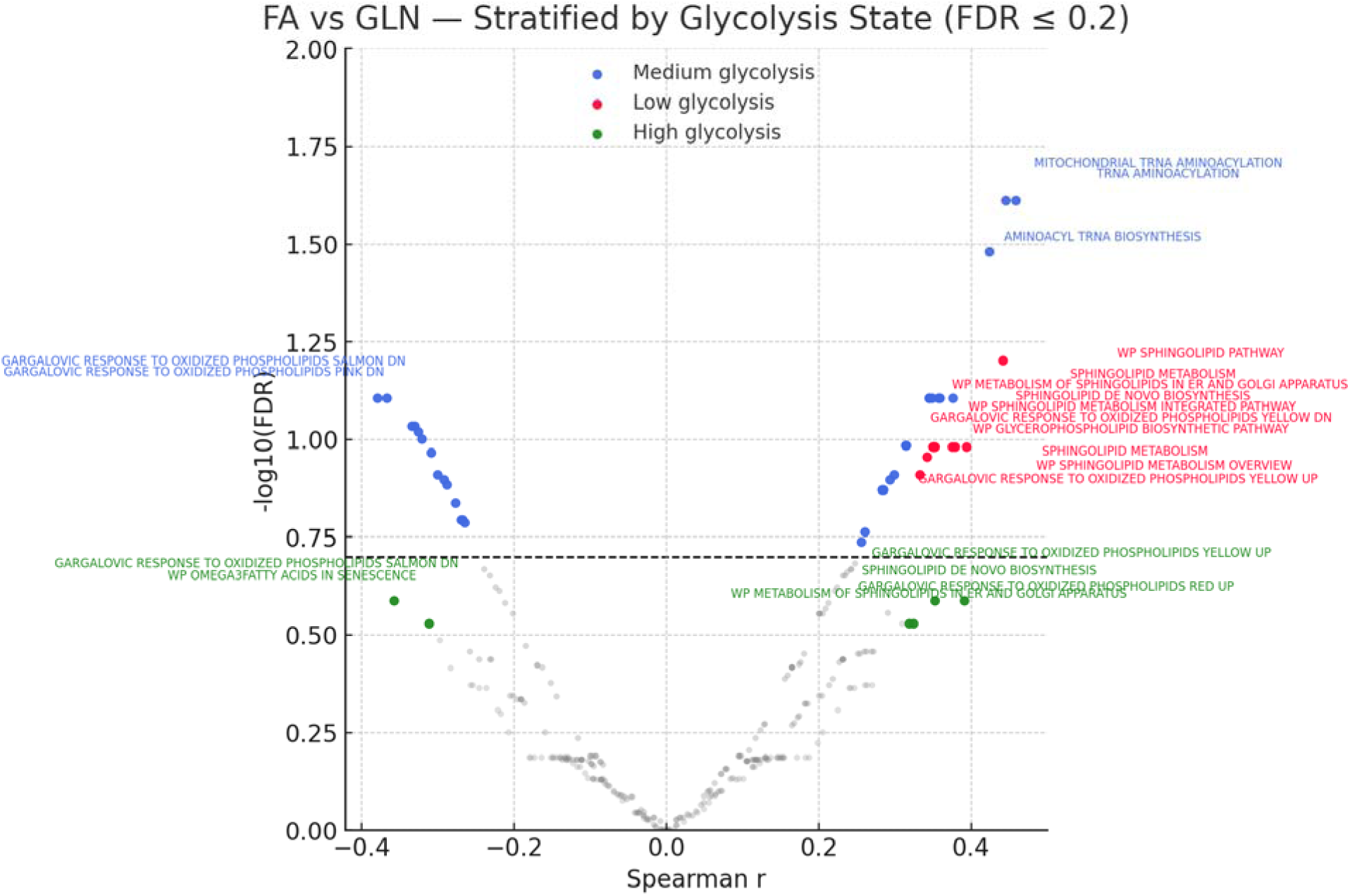
Volcano plots of FA-GLN correlations by glycolysis state. Spearman correlation coefficients (*ρ*) are shown on the x-axis and −log□□(FDR) on the y-axis. Points are colored by glycolysis state (Low = red, Medium = blue, High = green; approximately 57-58 tumors per group, n=172 total). Dashed lines indicate |ρ| = 0.20 and FDR ≤ 0.20 (significance). In Low glycolysis, several sphingolipid modules reached significance; in Medium glycolysis, aminoacyl-tRNA pathways reached significance. No pathways met FDR ≤ 0.20 in High glycolysis; near-threshold trends are displayed for context. *Abbreviations*: FA, fatty acid; GLN, glutamine; FDR, false discovery rate.

**Table 1.** Top fatty acid pathways correlated with glutamine activity by glycolytic state.

| Pathway (source) | Glycolysis state | Spearman $\rho$ | FDR (q) |
| --- | --- | --- | --- |
| Sphingolipid pathway (WikiPathways) | Low | 0.44 | 0.063 |
| Sphingolipid metabolism overview (WikiPathways) | Low | 0.39 | 0.104 |
| Sphingolipid metabolism (KEGG) | Low | 0.38 | 0.104 |
| Metabolism of sphingolipids in ER and Golgi apparatus (Reactome) | Low | 0.37 | 0.104 |
| Sphingolipid de novo biosynthesis (Reactome) | Low | 0.35 | 0.104 |
| Sphingolipid metabolism—integrated pathway (WikiPathways) | Low | 0.35 | 0.104 |
| Gargalovic response to oxidized phospholipids (MSigDB CGP) | Low | 0.35 | 0.104 |
| Glycerophospholipid biosynthetic pathway (Reactome/WikiPathways) | Low | 0.35 | 0.104 |
| Sphingolipid metabolism (Reactome) | Low | 0.34 | 0.111 |
| Gargalovic response to oxidized phospholipids (MSigDB CGP) | Low | 0.33 | 0.123 |
| tRNA aminoacylation (Reactome) | Medium | 0.46 | 0.024 |
| Mitochondrial tRNA aminoacylation (Reactome) | Medium | 0.45 | 0.024 |
| Aminoacyl-tRNA biosynthesis (KEGG) | Medium | 0.42 | 0.033 |
| Peroxisomal lipid metabolism (Reactome) | Medium | 0.38 | 0.078 |
| Mitochondrial fatty acid oxidation disorders (Reactome) | Medium | 0.36 | 0.078 |

*Abbreviations:* FDR, false discovery rate; ER, endoplasmic reticulum; MSigDB CGP, Molecular Signatures Database-chemical/genetic perturbations collection.

Elastic-Net regression did not outperform permutation-derived null distributions, indicating that FA pathways, while associated with GLN, do not provide strong predictive power for GLN activity. These aminoacyl-tRNA correlations are interpreted as reflecting general GLN-dependent biosynthetic demand rather than fatty acid-specific metabolic regulation.

### FA pathway activity and tumor stage

Linear regression models tested associations between FA pathway activity and AJCC stage (Fig 2, Table 2). Pathway activity generally decreased with advancing stage. The most negative coefficients were observed for Elevated circulating long-chain fatty acid concentrations (β_stage = –1.23, p = 0.183, FDR = 0.842), Very-long-chain fatty acid accumulation (β_stage = –0.92, p = 0.106, FDR = 0.981), Medium chain fatty-acyl-CoA metabolic process (β_stage = –0.86, p = 0.234, FDR = 0.697), and Very-long-chain fatty acid CoA ligase activity (β_stage = –0.74, p = 0.171, FDR = 0.981). None remained significant after multiple-testing correction.

**Fig 2.**
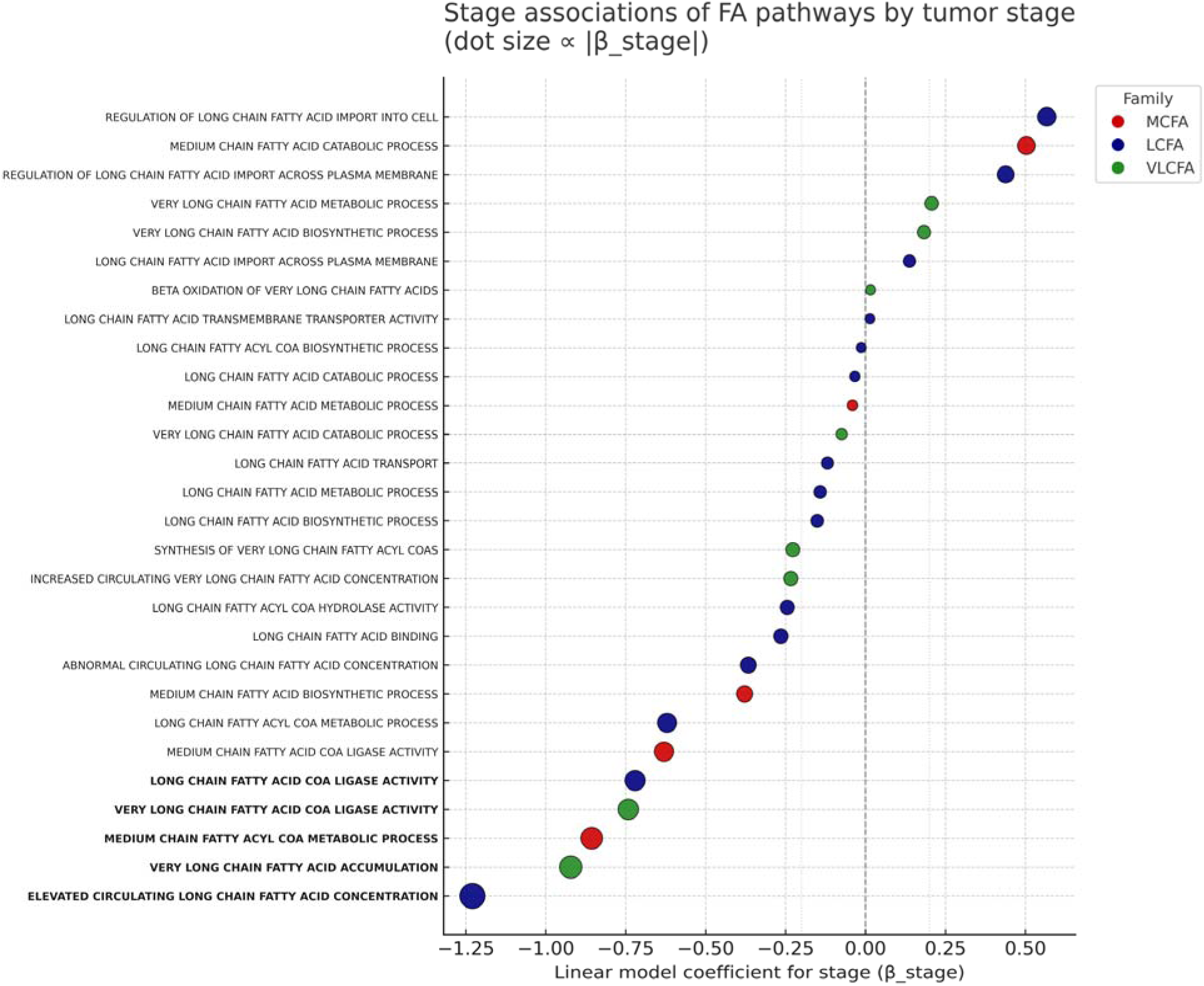
Associations between FA pathway activity and tumor stage. Multivariable linear regression coefficients for tumor stage (β_stage) are shown for FA pathways, adjusted for age, sex, and tumor mutational burden. Colors denote FA families (MCFA=red, LCFA=blue, VLCFA=green). The vertical dashed line marks no association (β_stage = 0). Pathways with the largest absolute β_stage within each family are highlighted. No pathway met FDR ≤ 0.25. Analyses were restricted to patients with complete clinical data (n = 166). *Abbreviations*: MCFA, medium chain fatty acid; LCFA, long chain fatty acid; VLCFA, very long chain fatty acid; TMB, tumor mutational burden.

**Table 2.** Associations between fatty acid pathway activity and tumor stage in PDAC. Regression coefficients for tumor stage (β_stage) were estimated from multivariable linear models of pathway activity, adjusted for age, sex, and tumor mutational burden (TMB). Pathways with |β_stage| ≥ 0.70 are shown (n = 166 patients with complete data). No pathway met the prespecified significance threshold of FDR ≤ 0.25.

| FA family | Pathway | $\beta_{\text{stage}}$ | P value | FDR (q) | R<sup>2</sup> |
| --- | --- | --- | --- | --- | --- |
| LCFA | Elevated circulating long-chain fatty acid concentrations | -1.23 | 0.183 | 0.842 | 0.015 |
| LCFA | Long chain fatty acid-CoA ligase activity | -0.72 | 0.116 | 0.842 | 0.024 |
| MCFA | Medium chain fatty-acyl-CoA metabolic process | -0.86 | 0.234 | 0.697 | 0.010 |
| VLCFA | Very long chain fatty acid accumulation | -0.92 | 0.106 | 0.981 | 0.018 |
| VLCFA | Very long chain fatty acid-CoA ligase activity | -0.74 | 0.171 | 0.981 | 0.016 |

*Abbreviations:* FA, fatty acid; LCFA, long chain FA; MCFA, medium chain FA; VLCFA, very long chain FA; TMB, tumor mutational burden; FDR, false discovery rate.
*Notes:* AJCC stage was treated as an ordinal variable (I-IV); $\beta_{\text{stage}}$ reflects the change in pathway activity per one-stage increase (negative values indicate lower activity at higher stage).

### Enzyme-class correlations with glutamine

Class-level meta correlations between FA enzyme length classes and GLN activity were estimated by pooling enzyme-specific Spearman correlations (Fisher’s z random-effects) and are shown in Fig 3 and Table 3. In glycolysis-medium tumors, very-long-chain fatty acid (VLCFA) enzymes showed the strongest positive correlation with GLN (r =0.23, 95% CI 0.09-0.34), followed by medium-chain fatty acid (MCFA) enzymes (r = 0.17, 95% CI 0.06-0.28). In glycolysis-low tumors, MCFA enzymes were weakly positive (r = 0.09, 95% CI –0.02 to 0.20), whereas VLCFA enzymes were near zero (r = 0.04, 95% CI –0.15 to 0.15). LCFA enzymes were weakly positive across states (r = 0.04-0.06). In glycolysis-high tumors, MCFA enzymes trended slightly negative (r = –0.05, 95% CI –0.20 to 0.10).

**Fig 3.**
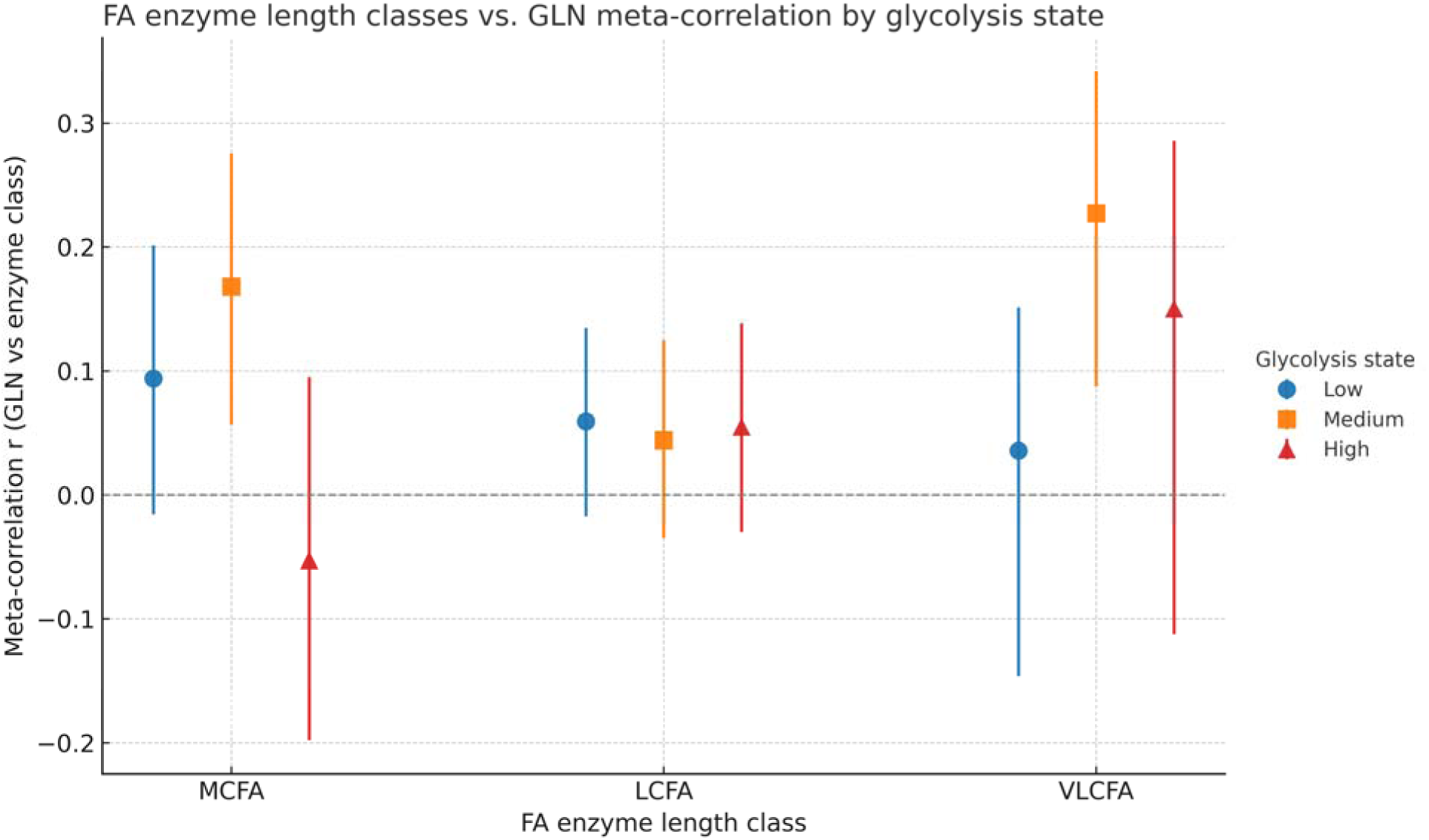
Association of FA enzyme length classes with GLN correlations across glycolysis states. Fisher’s meta-correlation coefficients (r) between enzyme-level FA activity and GLN activity are shown for MCFA, LCFA, and VLCFA classes within each glycolysis state (Low = blue, Medium = orange, High = red). Error bars indicate 95% confidence intervals from bootstrap resampling (B = 5,000); glycolysis strata are the same tertile groups used in Fig 1 (≈ 57-58 tumors per group). The vertical dashed line marks no association (r = 0). *Abbreviations*: MCFA, medium chain fatty acid; LCFA, long chain fatty acid; VLCFA, very long chain fatty acid.

**Table 3.**
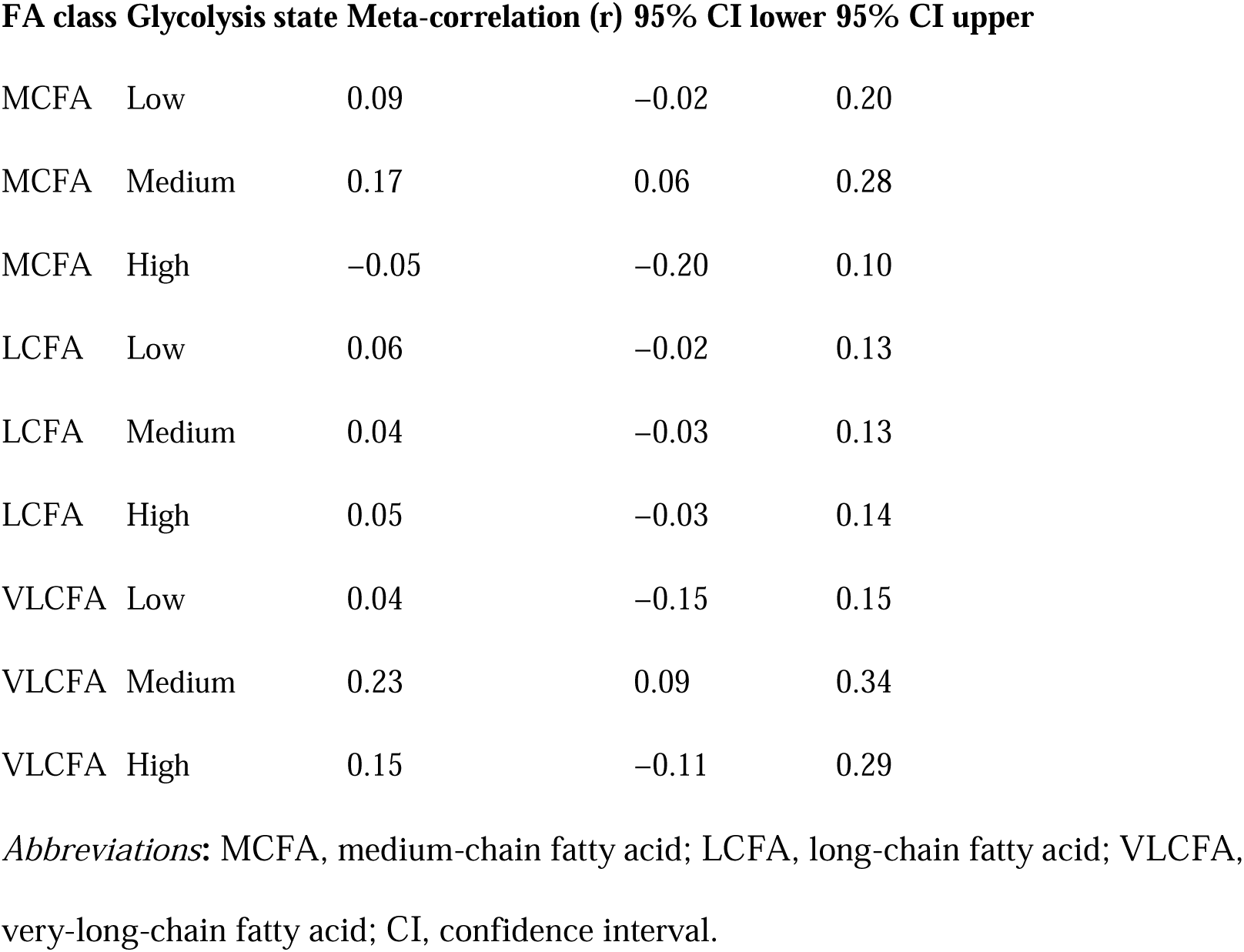
Meta-correlation of fatty acid enzyme classes with glutamine activity by glycolysis state. Fisher’s meta-correlation coefficients (r) were calculated within each glycolysis state for medium chain (MCFA), long chain (LCFA), and very long chain (VLCFA) enzyme classes. Values are shown with 95% confidence intervals from bootstrap resampling (B = 5,000). No associations met the prespecified FDR ≤ 0.25 threshold.

### Family-level comparisons

Family-level FA-GLN associations (Fig 4, Table 4; Supplementary Table S2 for family mappings) showed significant decreases in high versus low-glycolysis tumors for glycerophospholipids (Δr_meta = – 0.06, p = 0.0041, q = 0.021) and sphingolipids (Δr_meta = –0.12, p = 0.0019, q = 0.019). Other families (FA biosynthesis/elongation, FAO β-oxidation, lipid transport, triglycerides, unsaturated FA, FAO carnitine shuttle, lipid regulatory) did not pass q < 0.05.

**Fig 4.**
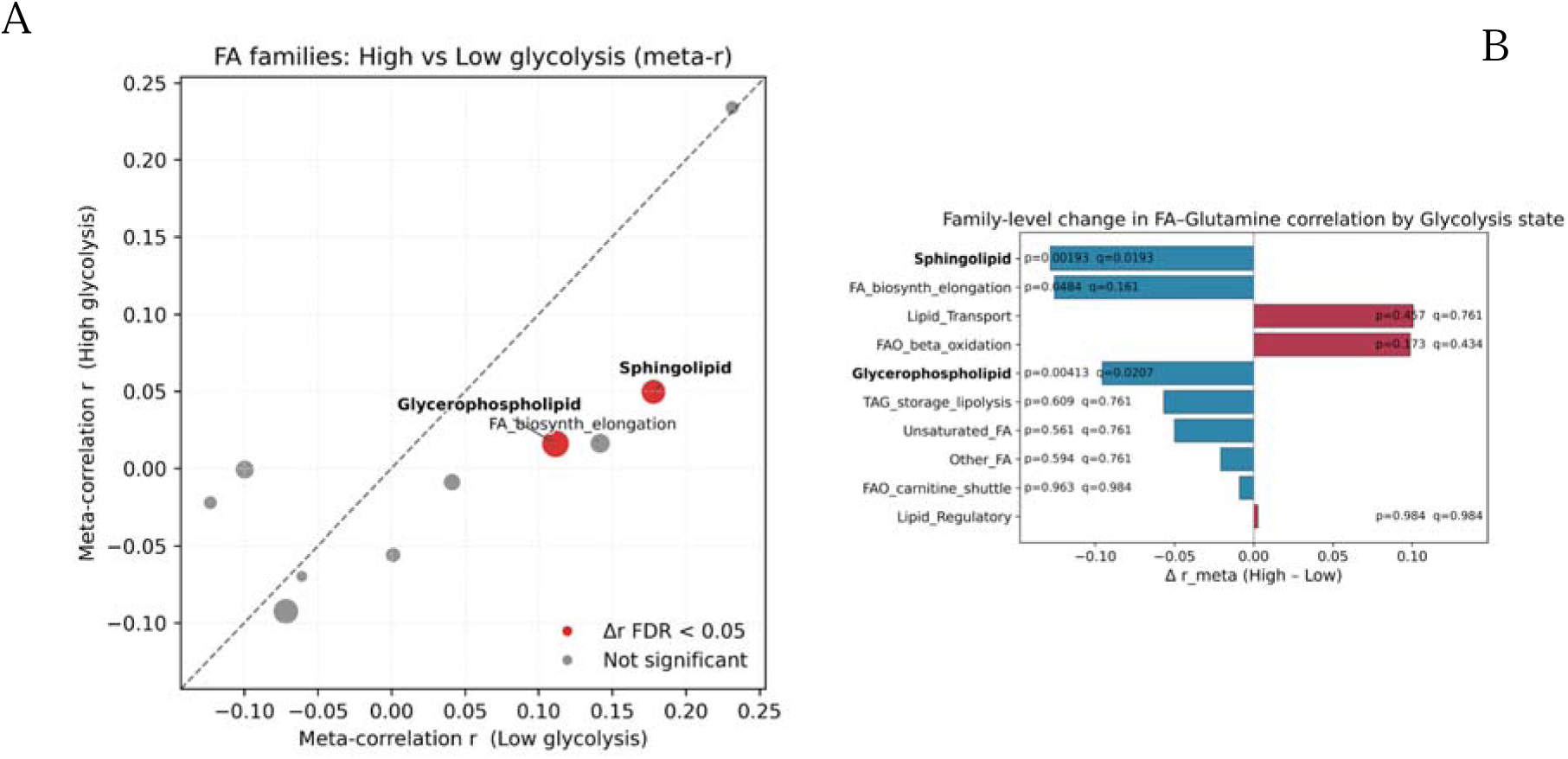
Family-level shifts in FA-GLN coupling by glycolysis state. (A) Scatterplot comparing family-level meta-correlations (r) in High versus Low glycolysis tumors; each point is an FA family. Families with significant differences (Δr, High-Low) at FDR q < 0.05 are highlighted. (B) Bar plot of Δr (High-Low) with corresponding p– and q-values; negative Δr indicates stronger FA-GLN coupling in Low glycolysis. Glycerophospholipid and sphingolipid families show significantly lower coupling in High versus Low glycolysis. *Abbreviations*: Δr, difference in meta-correlation; q, FDR-adjusted p-value.

**Table 4.**
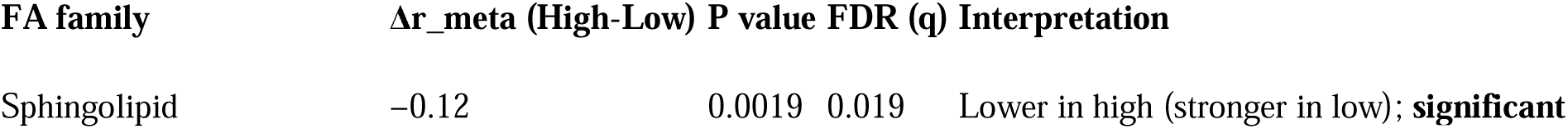
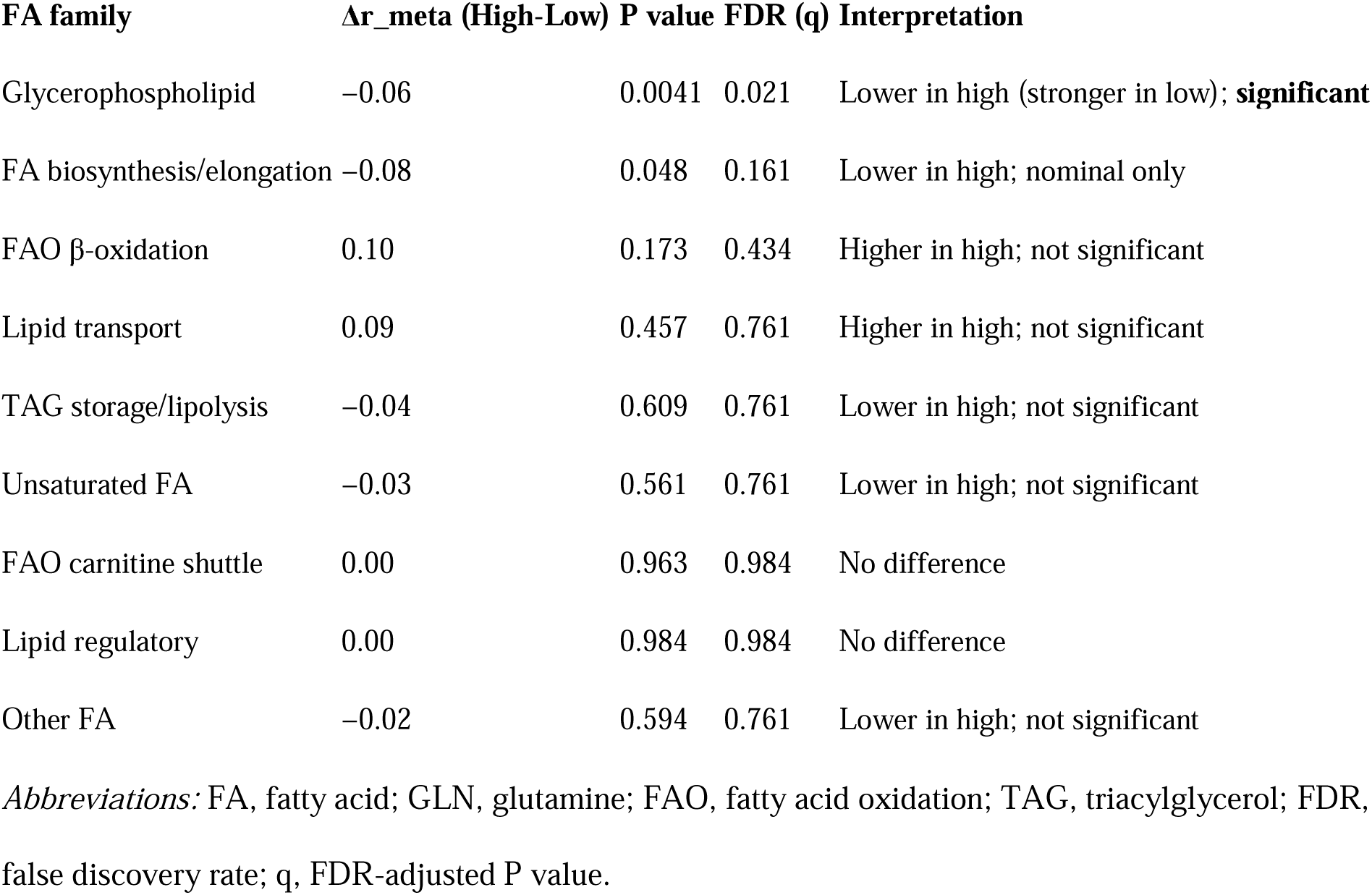
Family-level differences in fatty acid-glutamine associations by glycolysis state. Shown are Fisher-z meta-correlation differences (Δr, High-Low) for fatty acid (FA) families with two-sided P values and FDR-adjusted q values. Negative Δr indicates stronger FA-GLN coupling in glycolysis-low tumors; positive Δr indicates stronger coupling in glycolysis-high tumors. Family definitions/mappings are provided in S2 Table. FA family Δr_meta (High-Low).

### Survival analysis stratified by glycolysis state

Cox models were applied at enzyme-class and family levels (Fig 5; Tables 5,6). At the class level, LCFA activity was protective in glycolysis-high tumors (HR = 0.85, 95% CI 0.79-0.91, p = 1.0×10[_[_), borderline protective in glycolysis-medium tumors (HR = 0.93, 95% CI 0.86-1.01, p = 0.080), and not associated with survival in glycolysis-low tumors (HR = 0.99, 95% CI 0.93-1.06, p = 0.814). In glycolysis-low tumors, MCFA showed a borderline protective trend (HR = 0.89, 95% CI 0.78-1.02, p = 0.100). VLCFA trended toward higher hazard in glycolysis-medium tumors (HR = 1.10, 95% CI 0.98-1.23, p = 0.094). At the family level, in glycolysis-high tumors glycerophospholipid activity was associated with higher hazard (HR = 1.57, 95% CI 1.06-2.34, p = 0.025), whereas lipid transport (HR = 0.50, 95% CI 0.31-0.82, p = 0.006) and FAO carnitine shuttle (HR = 0.54, 95% CI 0.36-0.81, p = 0.003) were protective. In glycolysis-low tumors, lipid transport was also protective (HR= 0.64, 95% CI 0.42-0.99, p = 0.045). No families reached significance in the medium-glycolysis group.

**Fig 5.**
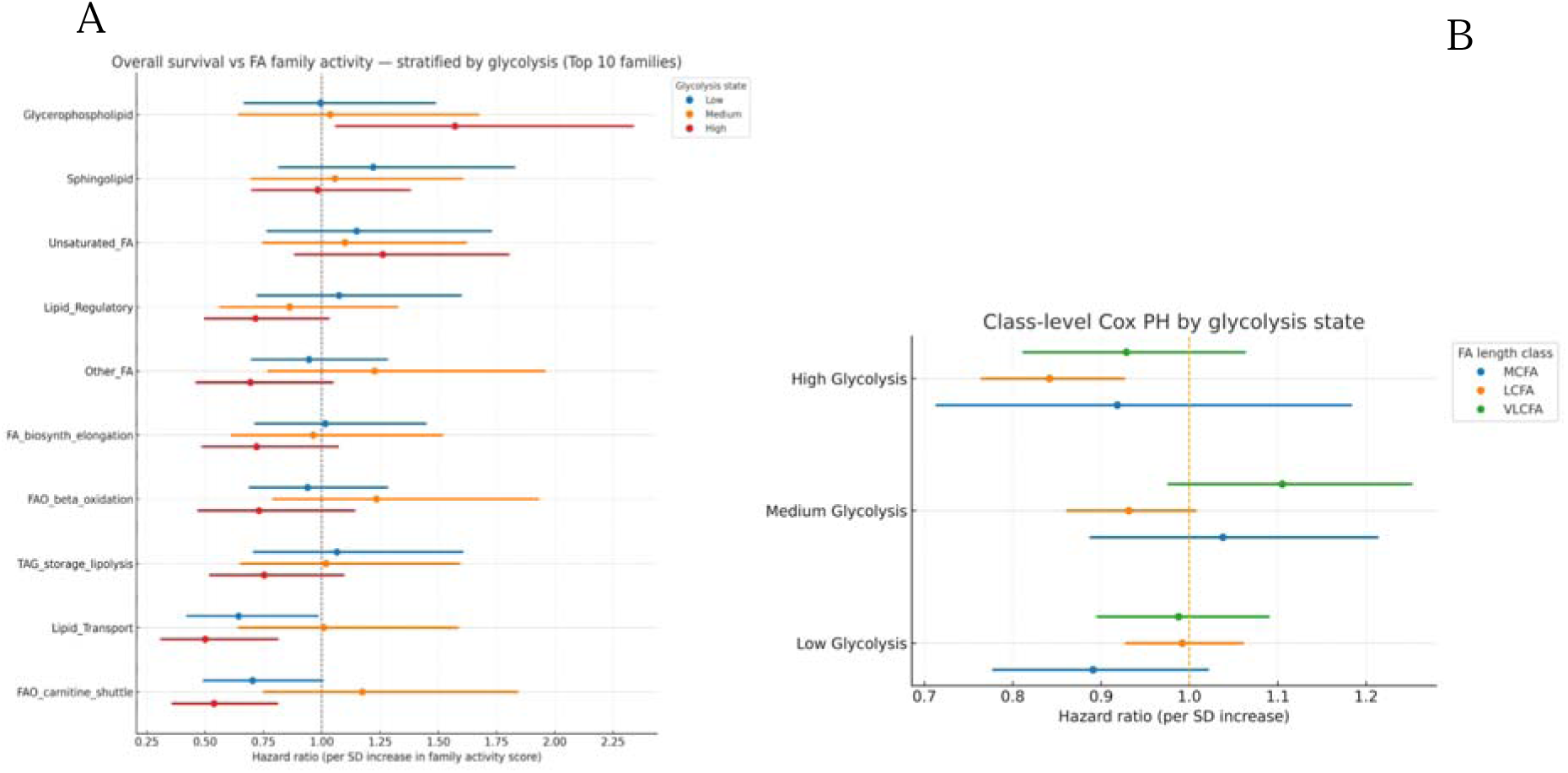
Survival associations of FA metabolism by family and chain length, stratified by glycolysis. Hazard ratios (HRs) for overall survival with 95% confidence intervals are shown (per 1-SD increase in activity), with models adjusted for age, sex, and tumor stage. (A) Top FA families, colors indicate glycolysis state (Low = blue, Medium = orange, High = red). (B) Enzyme classes (MCFA = blue, LCFA = orange, VLCFA = green). Models were fit separately within each glycolysis tertile group (Low, Medium, High), using the same stratification as in Fig 1. *Abbreviations*: MCFA, medium chain fatty acid; LCFA, long chain fatty acid; VLCFA, very long chain fatty acid.

**Table 5.**
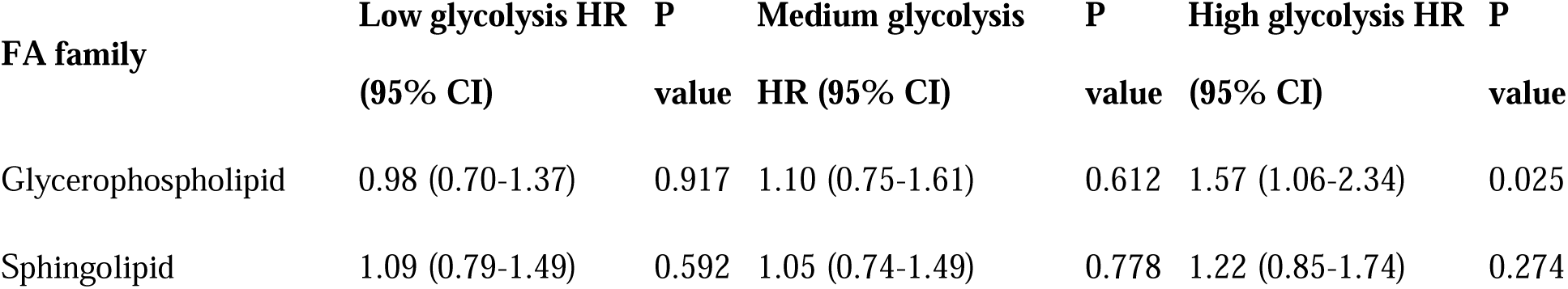
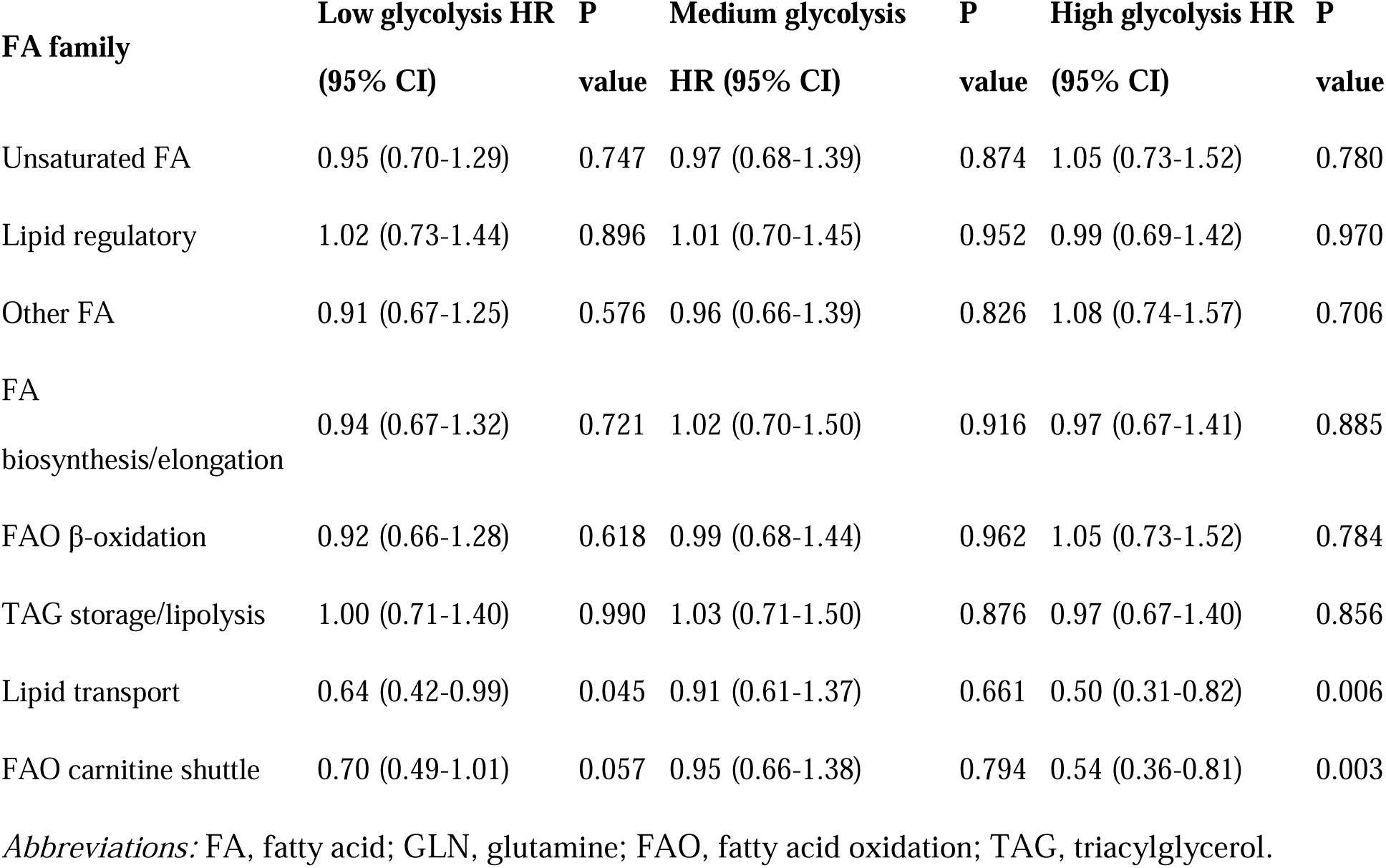
Family-level Cox proportional hazards models stratified by glycolysis state. Hazard ratios (HRs) are per 1-SD increase in family activity scores (single-sample gene set scoring–derived) and are adjusted for age, sex, and AJCC stage. P values are nominal (no FDR applied to survival models).

**Table 6.**
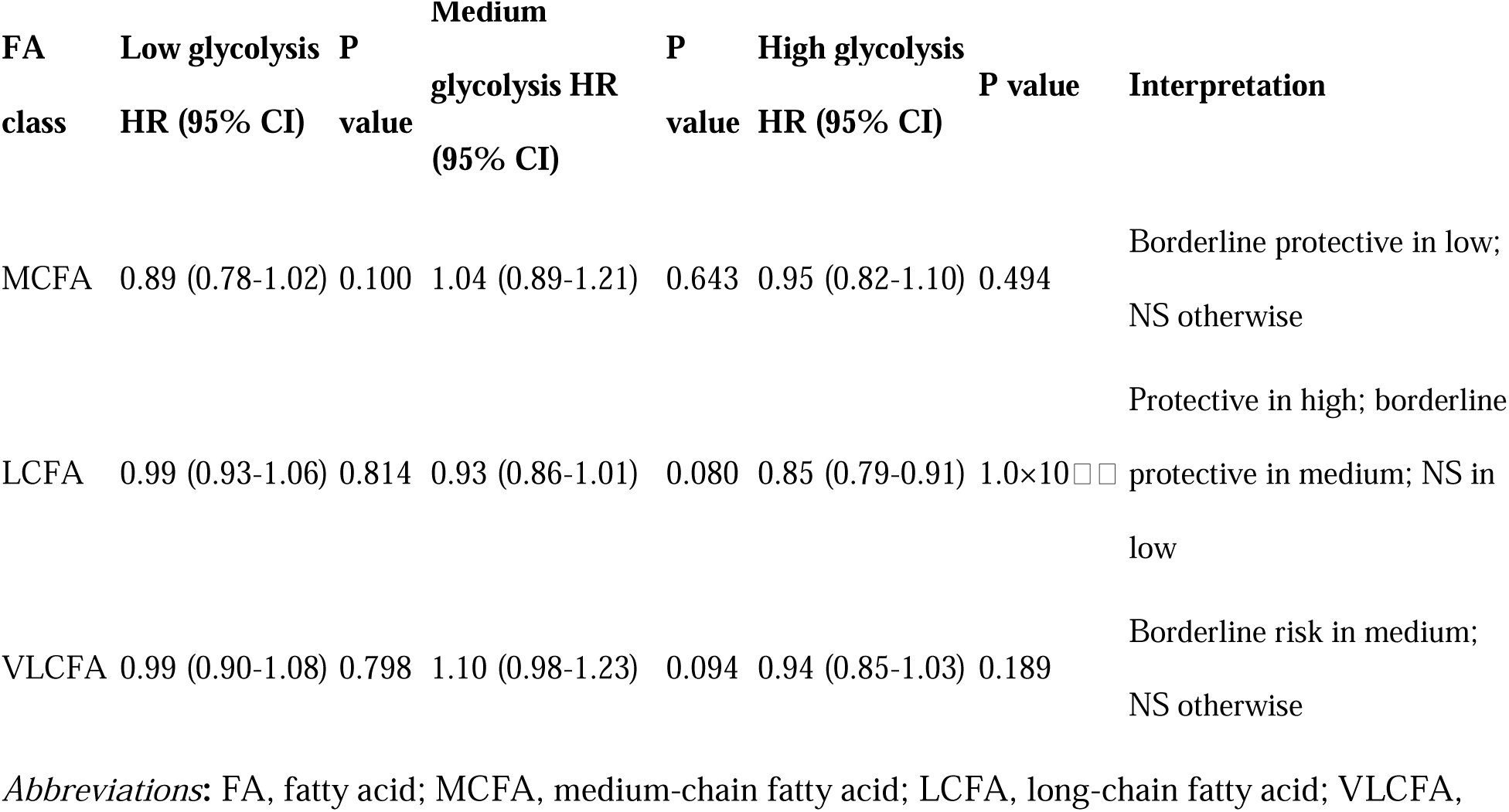
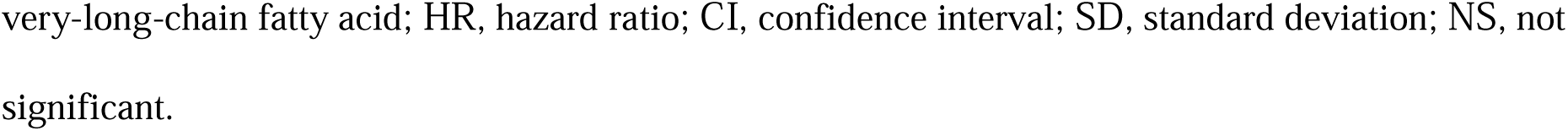
Class-level Cox proportional hazards models stratified by glycolysis state. Hazard ratios (HRs) are per 1-SD increase in class activity (MCFA, LCFA, VLCFA) and are adjusted for age, sex, and AJCC stage. P values are nominal (no FDR applied).

| FA<br>class | Low glycolysis<br>HR (95% CI) | P<br>value | Medium<br>glycolysis HR<br>(95% CI) | P<br>value | High glycolysis<br>HR (95% CI) | P value | Interpretation |
| --- | --- | --- | --- | --- | --- | --- | --- |
| MCFA | 0.89 (0.78-1.02) | 0.100 | 1.04 (0.89-1.21) | 0.643 | 0.95 (0.82-1.10) | 0.494 | Borderline protective in low;<br>NS otherwise |
| LCFA | 0.99 (0.93-1.06) | 0.814 | 0.93 (0.86-1.01) | 0.080 | 0.85 (0.79-0.91) | $1.0 \times 10^{-4}$ | Protective in high; borderline<br>protective in medium; NS in<br>low |
| VLCFA | 0.99 (0.90-1.08) | 0.798 | 1.10 (0.98-1.23) | 0.094 | 0.94 (0.85-1.03) | 0.189 | Borderline risk in medium;<br>NS otherwise |

*Abbreviations:* FA, fatty acid; MCFA, medium-chain fatty acid; LCFA, long-chain fatty acid; VLCFA,
very-long-chain fatty acid; HR, hazard ratio; CI, confidence interval; SD, standard deviation; NS, not significant.

### Supplementary predictive modeling

Permutation-based Elastic Net regression did not yield predictive performance beyond chance (Supplementary Fig S1, Supplementary Data S1, Supplementary Table S1): observed R² values did not differ significantly from the permutation-derived null distribution (1,000 permutations).

### Supplementary metabolite-inference validation of FA-GLN coordination

Metabolite activities inferred from transcriptomic data were evaluated across the 173 PDAC tumors in the GENIE cohort (Supplementary Fig. S3 and Supplementary Fig. S4; Supplementary Table S3; Supplementary Table S5; Supplementary Data S5). Thirteen metabolites representing glutamine metabolism (GLN, GLU, αKG), tricarboxylic acid (TCA) cycle intermediates (CITRATE, SUCCINATE, FUMARATE, MALATE), pyruvate-lactate exchange (PYRUVATE, LACTATE), fatty-acid oxidation (short/medium and long chain acyl-carnitines), very long chain fatty acid elongation (VLCFA_ELONG), and cellular redox buffering (GSH/GSSG) were analyzed. Unsupervised clustering of the resulting 13 × 173 activity matrix identified three metabolite-defined tumor states (Oxidative GLN-Driven, Hybrid FAO-TCA, Glycolytic FAO-Low), consistent with the dominant signatures illustrated in Supplementary Fig. S3 and Supplementary Fig. S4. A significant association was observed between glycolysis strata and the three metabolite-defined states (χ² = 14.54, df = 4, p = 0.0059; Cramér’s V = 0.21), indicating that coordinated metabolite phenotypes were modulated by glycolytic background. Differences in individual metabolite activities across the three states were examined using one-way ANOVA and the Kruskal-Wallis test. After Benjamini-Hochberg correction, statistically significant differences (FDR < 0.05) were detected for all 13 metabolites across the three states (Supplementary Table S4). The largest effects were observed for TCA-cycle intermediates (MALATE, SUCCINATE, FUMARATE), αKG, long chain acyl-carnitines, and VLCFA elongation, whereas PYRUVATE exhibited the weakest but still significant difference. Alignment between metabolite-defined states and transcriptomic FA-GLN coordination was examined. Low-glycolysis tumors, which previously showed the strongest GLN-TCA-lipid coordination transcriptomically, corresponded to the Oxidative GLN-Driven State. Medium-glycolysis tumors, which showed the strongest VLCFA-GLN coupling, corresponded to the Hybrid FAO-TCA State, characterized by elevated long-chain acyl-carnitines and increased FAO. High-glycolysis tumors, which demonstrated weak FA-GLN coordination, corresponded to the Glycolytic FAO-Low State, consistent with reduced acyl-carnitines and diminished oxidative metabolism. Together, these findings demonstrated that the FA-GLN metabolic hierarchy identified in pathway-level transcriptomic analyses was preserved when evaluated through metabolite-inference modeling and remained stratified by glycolytic background.

### Proteomic validation of FA-GLN family-level coordination

Protein-level analysis in the CPTAC PDAC cohort (n = 140 tumors) provided an orthogonal test of family-level FA-GLN coordination (S7 Fig, S7 Table). Using Hallmark Glycolysis protein-module scores, tumors were divided into low, medium, and high glycolysis tertiles (47, 46, and 47 tumors, respectively), and GLN, glycerophospholipid (GPL), and sphingolipid (SP) protein modules were scored from the curated gene lists (S2 Table) after filtering for proteins with at least 70% observed values.

Sphingolipid-GLN coordination was strongest in low– and medium-glycolysis tumors and attenuated in high-glycolysis tumors. The SP protein module exhibited a moderate negative correlation with GLN in low glycolysis (Spearman r = –0.36, q = 0.036) and in medium glycolysis (r = –0.45, q = 0.011), but was near zero in high glycolysis (r = 0.04, q = 0.93). GPL protein scores showed small, non-significant correlations with GLN in all strata (|r| ≤ 0.17, q ≥ 0.52). When changes were summarized as differences in correlation magnitude between high and low glycolysis, sphingolipids showed a pronounced loss of GLN coupling in high-glycolysis tumors, whereas glycerophospholipids showed minimal change.

Although the direction of the SP-GLN correlation differed between RNA and protein layers (positive in the transcriptomic pathway analysis and negative in the proteomic analysis), both datasets supported the conclusion that sphingolipid programs are preferentially linked to GLN metabolism in tumors with lower glycolytic activity and become uncoupled as glycolysis increases.

## Discussion

This study profiled fatty acid (FA) metabolism in PDAC patient tumors with attention to chain length, lipid families, and glycolytic state. Three consistent findings emerged. First, FA-GLN coupling was strongly context-dependent: in glycolysis-low tumors, sphingolipid and glycerophospholipid pathways correlated most strongly with GLN activity, whereas aminoacyl-tRNA pathways dominated in glycolysis-medium tumors, and few FA pathways correlated with GLN in glycolysis-high tumors. Second, enzyme-class meta-analysis indicated that VLCFA enzymes showed the tightest positive association with GLN in medium-glycolysis tumors, while MCFA enzymes weakened or inverted their association in glycolysis-high tumors. Third, family-level correlations for glycerophospholipids and sphingolipids were significantly lower in high versus low glycolysis tumors, suggesting a decoupling of lipid remodeling from GLN metabolism as glycolytic dependence increases.

Clinical associations paralleled these patterns. At the class level, LCFA activity was protective in glycolysis-high tumors, borderline protective in glycolysis-medium tumors, and not associated with outcome in glycolysis-low tumors. At the family level, glycerophospholipid activity associated with higher hazard in glycolysis-high tumors, whereas lipid transport and FAO carnitine-shuttle activity were protective in both high and low glycolysis states. Together, these results argue that FA biology in PDAC is heterogeneous and that prognostic effects depend on both chain length and glycolytic context.

These observations provide a patient-based complement to preclinical work showing that altering nutrient availability (for example, ketogenic-diet conditions) modulates responsiveness to GLN-pathway inhibition. The present data refine that concept by indicating that not all FA classes behave similarly across glycolytic states: membrane-focused families (glycerophospholipids, sphingolipids) appear preferentially coupled to GLN in glycolysis-low tumors, whereas in glycolysis-high tumors the same families lose this coupling and glycerophospholipid activity associates with worse survival. One interpretation is that when glycolysis is low, GLN anaplerosis and lipid remodeling are coordinated to sustain biomass and signaling; as glycolysis rises, carbohydrate-derived carbons may partially substitute for GLN-linked lipid programs, weakening coupling and shifting the prognostic role of specific FA families. The stronger GLN association for VLCFA enzymes in medium-glycolysis tumors may reflect increased demands for membrane elongation/desaturation or peroxisomal contributions during intermediate metabolic states. These mechanisms remain hypotheses that require flux-level validation.

Additionally, Elastic-Net analysis confirmed that FA pathways, despite their associations with GLN activity, do not provide strong predictive power for GLN levels, indicating that these relationships reflect coordinated metabolic patterns rather than direct predictiveness. In medium-glycolysis tumors, the prominence of aminoacyl-tRNA pathways is interpreted as reflecting GLN-dependent biosynthetic activity rather than FA-specific coordination.

To evaluate whether these transcriptomically derived FA-GLN coordination patterns reflected broader metabolic programs, a supplementary metabolite-inference analysis was performed. Three metabolite-defined states were identified and were shown to differ significantly across all evaluated metabolites. These states paralleled the FA-GLN hierarchy: the Oxidative GLN-Driven State corresponded to low-glycolysis tumors, the Hybrid FAO-TCA State corresponded to medium-glycolysis tumors, and the Glycolytic FAO-Low State corresponded to high-glycolysis tumors. The preservation of these relationships indicated that the FA-GLN metabolic architecture observed in pathway-level transcriptomic analyses reflected broader, coordinated metabolic programs when evaluated from a metabolite-inference perspective.

Independent protein-level validation in the CPTAC PDAC cohort supported the glycolysis-stratified FA-GLN hierarchy observed at the transcriptomic and metabolite-inference levels. Sphingolipid protein modules showed stronger GLN coupling in low– and medium-glycolysis tumors than in high-glycolysis tumors, whereas glycerophospholipid protein modules exhibited only weak, non-significant associations with GLN. Although the sign of the sphingolipid-GLN correlation differed between the transcriptomic and proteomic analyses, both layers converged on a model in which sphingolipid programs are preferentially coupled to GLN metabolism in less glycolytic tumors and become attenuated as glycolytic dependence increases. This multi-layer agreement, together with the metabolite-inference results, suggests that the observed FA-GLN relationships reflect broader coordinated metabolic programs rather than cohort-specific artifacts. Limitations of this study include reliance on bulk RNA-seq and enrichment-based pathway activity (which do not measure flux or resolve cell-type contributions) and reduced power within glycolysis strata for survival analyses, where events are further partitioned across the three glycolysis groups.

Glycolysis tertiles were selected a priori for stratification, and alternative cut-points were not systematically explored, so some findings may depend on this choice. Gene-set definitions of FA “families” may not capture all biochemical specificities, and results may be sensitive to curation choices. The study is observational and cross-sectional, so causal inferences cannot be drawn. Finally, predictive modeling with Elastic Net did not outperform permutation nulls, indicating that while associations are robust, GLN activity is not readily predicted from FA pathways alone within strata.

### Implications and next steps

Stratifying PDAC by glycolytic state may be necessary when evaluating FA or GLN-directed interventions and when interpreting dietary manipulations that shift circulating FA composition. Prospective studies should (i) combine tumor glycolysis readouts (for example, transcriptomic scores or FDG-PET) with lipidomics and stable-isotope tracing, (ii) validate chain length-specific effects in controlled perturbations (MCFA versus LCFA availability) with and without GLN pathway inhibitors, and (iii) use single-cell and spatial approaches to resolve tumor-stroma contributions to FA and GLN programs.

## Conclusion

FA metabolism in PDAC is not uniform; its associations with GLN metabolism and with patient outcomes depend on both chain length and glycolytic state. Glycerophospholipid and sphingolipid families show the strongest link to GLN in glycolysis-low tumors but are decoupled in glycolysis-high tumors, where glycerophospholipid activity associates with poorer survival. LCFA activity is protective in glycolysis-high tumors and shows a borderline protective trend in medium glycolysis. These glycolysis stratified patterns define conditional metabolic vulnerabilities and provide a rationale to incorporate tumor glycolytic state and FA class composition when designing GLN-targeted or diet-modulated therapeutic strategies.

## Methods

### Data source and preprocessing

RNA-seq and clinical data from the AACR Project GENIE PDAC cohort were analyzed [20]. Clinical variables included age, sex, stage, and tumor mutational burden (TMB). Gene-level expression was quantified as log[_ transcripts per million (TPM). Quality control included verification of identifiers, consistency of clinical annotations, and exclusion of samples failing RNA-seq metrics (e.g., low library complexity, missing covariates). Genes with low expression (counts per million < 1 in > 80% of samples) were excluded. One sample had complete expression data but lacked clinical annotations; therefore, analyses requiring clinical covariates used n=172, while metabolite-inference analyses used n=173.

### Pathway curation

FA and GLN pathways were curated from KEGG and Reactome [21,22], including biosynthesis and degradation processes. Pathways were grouped into MCFA, LCFA, and VLCFA classes to capture chain length-specific biology [14,17]. In addition to chain-length classes, pathways and enzymes were grouped into functional FA families (e.g., sphingolipids, glycerophospholipids, triglycerides, lipid transport, FA biosynthesis/elongation, FA oxidation, FAO carnitine shuttle, lipid regulatory, and unsaturated FA) using KEGG/Reactome and WikiPathways annotations [23]; families could span chain-length classes.

Glycolysis stratification used the MSigDB Hallmark Glycolysis gene set [24]. The curated mapping of pathways to fatty acid (FA) families is provided in Supplementary Table S2. A schematic showing representative enzymes associated with each chain-length class and lipid family is presented in Supplementary Fig. S6. When a FA pathway appeared in both a chain-length class and a lipid family (e.g., ELOVL1 in both LCFA and SP), we retained its membership in both categories during correlation and enrichment analyses without double-counting samples. Overlapping assignments were biologically meaningful and retained, as pathways may participate in more than one functional grouping. Each grouping was analyzed independently, allowing shared pathways to contribute to multiple interpretations.

### Pathway activity scoring

FA and GLN pathway activity scores were computed using gene set variation analysis (GSVA) implemented in the GSVA R package [25]. Glycolysis scores used for stratification were computed using single-sample gene set enrichment analysis (ssGSEA) of the MSigDB Hallmark Glycolysis gene set [24], and tumors were divided into low, medium, and high tertiles (≈57–58 tumors per group; n=172 total). Pathway scores were z-standardized so that effects were expressed per one standard deviation (1 SD) increase.

### FA-GLN correlations stratified by glycolysis state

Spearman correlations were calculated between FA and GLN pathway activity within each glycolysis stratum. Multiple testing correction used the Benjamini-Hochberg false discovery rate (FDR) procedure with a threshold of FDR ≤ 0.20 [26].

### FA pathway activity and tumor stage

Associations between FA activity and AJCC stage were evaluated with linear regression adjusted for age, sex, and TMB. Regression coefficients (β_stage), p-values, FDR-adjusted q-values, and R² were reported.

### Enzyme-class correlations with glutamine

Enzyme activity was defined as the z-score of expression for genes encoding FA metabolic enzymes. Enzyme-GLN correlations were estimated within glycolysis strata. Correlations were aggregated into MCFA, LCFA, and VLCFA classes using random-effects meta-analysis with inverse-variance weighting of Fisher’s z-transformed values; pooled estimates were back-transformed to correlation coefficients. Bootstrap resampling (B = 5,000) was used to quantify uncertainty.

### Family-level comparisons

Enzyme-level correlations were combined within families using Fisher’s z-transformation to obtain pooled meta-correlations. Differences between glycolysis states (Δr_meta) were assessed with bootstrap resampling (B = 5,000). Two-sided p-values were derived from bootstrap distributions and FDR-adjusted across families.

### Survival analysis stratified by glycolysis

Overall survival was analyzed using Cox proportional hazards models adjusted for age, sex, and stage; hazard ratios (HRs) and 95% confidence intervals (CIs) were reported [27].

- Enzyme-class level: enzyme-specific log(HR) estimates were pooled within MCFA, LCFA, and VLCFA classes using random-effects meta-analysis.
- Family level: FA family scores (derived from single-sample gene set scoring) were evaluated within each glycolysis stratum using Cox models.

### Predictive modeling

Elastic Net regression tested whether FA activity predicted GLN activity within glycolysis strata. Models included age, sex, stage, and TMB as covariates. Performance was evaluated using R². Permutation testing (1,000 permutations) generated null R² distributions for comparison [28,29].

### Supplementary Metabolite Inference and Metabolic-State Assignment

A curated metabolite-gene mapping was assembled to infer activity for 13 metabolites relevant to glutamine metabolism (GLN, GLU, αKG), tricarboxylic acid (TCA) cycle intermediates (CITRATE, SUCCINATE, FUMARATE, MALATE), pyruvate-lactate exchange (PYRUVATE, LACTATE), fatty-acid oxidation (short/medium and long chain acyl-carnitines), very long chain fatty acid elongation (VLCFA_ELONG), and cellular redox buffering (GSH/GSSG). Enzymatic reactions were annotated as production or consumption steps, and associated genes were assigned weights of +1 or –1 accordingly. For each metabolite, a weighted mean expression value was calculated across all mapped genes. All metabolite scores were then z-normalized across samples to produce a 13 × 173 activity matrix. Principal component analysis followed by k = 3 clustering was applied to the normalized metabolite activity matrix to define three metabolite-based tumor states (Supplementary Fig. S3 and Supplementary Fig. S4). These states were annotated based on characteristic metabolic signatures and were designated as the Oxidative GLN-Driven State (GLN_TCA_REDOX_HIGH), the Hybrid FAO-TCA State (TCA_GLYCOLYTIC_FAO_HIGH), and the Glycolytic FAO-Low State (FAO_LOW). Associations between the three metabolite-defined states and the glycolysis strata (Low, Medium, High) were evaluated using the chi-square test of independence, and effect size was quantified using Cramér’s V. Differences in individual metabolite activities across the three metabolite-defined states were evaluated using one-way ANOVA and the Kruskal-Wallis test. All analyses were performed only when ≥2 samples per state were available, and statistical significance was determined using Benjamini-Hochberg FDR correction. Significant differences (FDR < 0.05) were detected for all 13 metabolites in both tests, confirming that each metabolite-defined state represented a distinct metabolic phenotype. Complete statistical outputs for metabolite activity comparisons are provided in Supplementary Table S4, and the metabolite-gene mapping and metabolic-state labels are provided in Supplementary Table S5 and Supplementary Data S5, respectively.

### Proteomic validation in the CPTAC PDAC cohort

Protein-level validation of fatty acid-glutamine (FA-GLN) relationships was performed using the CPTAC PDAC cohort described by Cao et al. [34]. Gene-level protein abundance data (log□-transformed, median-centered) were obtained from LinkedOmics (CPTAC-PDAC, MD abundance tumor matrix) and restricted to primary tumors with matched proteomic profiles, yielding 140 tumors with usable data. Descriptive statistics for the CPTAC proteomic cohort are provided in S6 Table.

Hallmark Glycolysis, GLN, glycerophospholipid (GPL), and sphingolipid (SP) modules were constructed by intersecting the curated transcriptomic gene sets (S2 Table) with the proteome and retaining proteins with at least 70% non-missing values across samples. For each module, protein scores were computed by z-scoring individual proteins across tumors and averaging z-scores within each module. Tumors were stratified into low, medium, and high glycolysis groups by tertiles of the Hallmark Glycolysis protein score (47, 46, and 47 tumors, respectively), mirroring the transcriptomic analysis.

Within each glycolysis tertile, Spearman correlations were calculated between GLN protein scores and each FA family protein score (GPL, SP). Two-sided p-values were adjusted for multiple testing using the Benjamini-Hochberg FDR procedure across all family-stratum combinations. Results are summarized in Supplementary Table S7 and visualized in Supplementary Fig. S7.

### Implementation

Analyses were performed in R (v4.3) and Python (v3.12). GSVA was run with the GSVA package [25]. Glycolysis scores used for stratification were computed using ssGSEA for the MSigDB Hallmark Glycolysis set [24]. In Python, data handling used NumPy and pandas [30,31]; Elastic Net and permutation tests were implemented with scikit-learn [32]; figures were generated with Matplotlib [33].

*Prespecified statistical thresholds differed by analysis type: FA-GLN pathway correlations used an FDR* ≤ *0.20; tumor-stage associations used FDR* ≤ *0.25 due to smaller effect sizes; analyses using q-values (family-level* Δ*r comparisons and metabolite-inference tests) used FDR < 0.05; and survival analyses report nominal p-values without FDR correction. The FDR* ≤ *0.20 threshold for pathway-level FA-GLN correlations was chosen as an exploratory screen to maintain sensitivity in the context of a moderate cohort size and a correlated set of pathways, whereas the principal family-level and metabolite-inference findings emphasized in the text satisfy stricter significance criteria (q < 0.05). Together, this hierarchical use of thresholds serves as a built-in sensitivity analysis, in which broad pathway screens are followed by more stringent family-level and metabolite-based tests before results are interpreted*.

## Data availability

All supplementary figures, tables, and source data used in this study are publicly available on Zenodo (DOI: 10.5281/zenodo.17870791). GENIE PDAC RNA-seq and associated clinical annotations are available through AACR Project GENIE. CPTAC-PDAC proteomics data are available through CPTAC/LinkedOmics as described in the original CPTAC PDAC publication. Code is available upon reasonable request.

### Ethics approval and consent to participate

This study used only de-identified, publicly available human tumor datasets from the AACR Project GENIE consortium and the Clinical Proteomic Tumor Analysis Consortium (CPTAC). All samples were collected under the oversight of the original studies’ institutional review boards, and informed consent was obtained by those studies. No new patient recruitment, intervention, or identifiable data were involved in the present analysis. In accordance with institutional and journal policies, additional ethics approval for secondary analysis of anonymized public data was not required.

### Consent for publication

Not applicable; this work analyzes de-identified public datasets and does not include individual-person data, images, or case descriptions requiring consent.

## References

1. Siegel RL, Miller KD, Fuchs HE, Jemal A. Cancer statistics, 2024. CA Cancer J Clin. 2024;74(1):17–48.

2. Rahib L, Smith BD, Aizenberg R, Rosenzweig AB, Fleshman JM, Matrisian LM. Projecting cancer incidence and deaths to 2030: the unexpected burden of thyroid, liver, and pancreas cancers. Cancer Res. 2014;74(11):2913–21.

3. Pavlova NN, Thompson CB. The emerging hallmarks of cancer metabolism. Cell Metab. 2016;23(1):27–47.

4. Vander Heiden MG, DeBerardinis RJ. Understanding the intersections between metabolism and cancer biology. Cell. 2017;168(4):657–69.

5. Hanahan D, Weinberg RA. Hallmarks of cancer: the next generation. Cell. 2011;144(5):646–74.

6. Kamphorst JJ, Nofal M, Commisso C, Hackett SR, Lu W, Grabocka E, et al. Human pancreatic tumors are nutrient poor and tumor cells actively scavenge extracellular protein. Cancer Res. 2015;75(3):544–53.

7. Ying H, Kimmelman AC, Lyssiotis CA, Hua S, Chu GC, Fletcher-Sananikone E, et al. Oncogenic KRAS maintains pancreatic tumors through regulation of anabolic glucose metabolism. Cell. 2012;149(3):656–70.

8. Bailey P, Chang DK, Nones K, Johns AL, Patch AM, Gingras MC, et al. Genomic analyses identify molecular subtypes of pancreatic cancer. Nature. 2016;531(7592):47–52.

9. Altman BJ, Stine ZE, Dang CV. From Krebs to clinic: glutamine metabolism to cancer therapy. Nat Rev Cancer. 2016;16(10):619–34.

10. Son J, Lyssiotis CA, Ying H, Wang X, Hua S, Ligorio M, et al. Glutamine supports pancreatic cancer growth through a KRAS-regulated metabolic pathway. Nature. 2013;496(7443):101–5.

11. Hensley CT, Wasti AT, DeBerardinis RJ. Glutamine and cancer: cell biology, physiology, and clinical opportunities. J Clin Invest. 2013;123(9):3678–84.

12. Encarnación-Rosado J, Chen HC, Tu J, Taniguchi K, Guo Q, Cui Y, et al. Pharmacological inhibition of glutamine metabolism by DON induces metabolic crisis in PDAC. Cancer Discov. 2024;14(5):1123–35.

13. Sousa CM, Biancur DE, Wang X, Halbrook CJ, Sherman MH, Zhang L, et al. Pancreatic stellate cells support tumor metabolism through autophagic alanine secretion. Nature. 2016;536(7617):479–83.

14. Röhrig F, Schulze A. The multifaceted roles of fatty acid synthesis in cancer. Nat Rev Cancer. 2016;16(11):732–49.

15. Koundouros N, Poulogiannis G. Reprogramming of fatty acid metabolism in cancer. Br J Cancer. 2020;122(1):4–22.

16. Schug ZT, Vande Voorde J, Gottlieb E. The metabolic fate of acetate in cancer. Nat Rev Cancer. 2016;16(11):708–17.

17. Nagarajan SR, Butler LM, Hoy AJ. The diversity and breadth of cancer cell fatty acid metabolism. Cancer Metab. 2021; 9:2.

18. Hajihassani O, Zarei M, Roichman A, Sadeh R, Eilam R, Wainberg Z, et al. A ketogenic diet sensitizes pancreatic cancer to inhibition of glutamine metabolism. bioRxiv. 2024. doi:10.1101/2024.07.19.604377.

19. Jin J, Byun J-K, Choi Y-K, Park K-G. Targeting glutamine metabolism as a therapeutic strategy for cancer. Exp Mol Med. 2023; 55:706–15.

20. AACR Project GENIE Consortium. AACR Project GENIE: powering precision medicine through an international consortium. Cancer Discov. 2017;7(8):818–31.

21. Kanehisa M, Furumichi M, Sato Y, Kawashima M, Ishiguro-Watanabe M. KEGG for taxonomy-based analysis of pathways and genomes. Nucleic Acids Res. 2023;51(D1): D587–92.

22. Jassal B, Matthews L, Viteri G, Gong C, Lorente P, Fabregat A, et al. The Reactome pathway knowledgebase. Nucleic Acids Res. 2020;48(D1): D498–503.

23. Kutmon M, Riutta A, Nunes N, Hanspers K, Willighagen EL, Bohler A, et al. WikiPathways: capturing the full diversity of pathway knowledge. Nucleic Acids Res. 2016;44(D1): D661–7.

24. Liberzon A, Birger C, Thorvaldsdóttir H, Ghandi M, Mesirov JP, Tamayo P. The MSigDB hallmark gene-set collection. Cell Syst. 2015;1(6):417–25.

25. Hänzelmann S, Castelo R, Guinney J. GSVA: gene set variation analysis for microarray and RNA-seq data. BMC Bioinformatics. 2013; 14:7.

26. Benjamini Y, Hochberg Y. Controlling the false discovery rate: a practical and powerful approach to multiple testing. J R Stat Soc B. 1995;57(1):289–300.

27. Cox DR. Regression models and life-tables. J R Stat Soc B. 1972;34(2):187–220.

28. Zou H, Hastie T. Regularization and variable selection via the elastic net. J R Stat Soc B. 2005;67(2):301–20.

29. Efron B, Tibshirani RJ. An Introduction to the Bootstrap. Boca Raton: Chapman & Hall/CRC; 1994.

30. Harris CR, Millman KJ, van der Walt SJ, Gommers R, Virtanen P, Cournapeau D, et al. Array programming with NumPy. Nature. 2020;585(7825):357–62.

31. McKinney, W. Data structures for statistical computing in Python. In: van der Walt S, Millman J, editors. Proceedings of the 9th Python in Science Conference. 2010. p. 51–56.

32. Pedregosa F, Varoquaux G, Gramfort A, Michel V, Thirion B, Grisel O, et al. Scikit-learn: Machine learning in Python. J Mach Learn Res. 2011; 12:2825–30.

33. Hunter JD. Matplotlib: A 2D graphics environment. Comput Sci Eng. 2007;9(3):90–5.

34. Cao L, Huang C, Cui Zhou D, Hu Y, Lih T-M, Pan J, et al. Proteogenomic characterization of pancreatic ductal adenocarcinoma. Cell. 2021;184(19):5031–5052.e26.

