## Supplementary material for "Glycolysis-stratified coordination of fatty acid and glutamine metabolism in pancreatic ductal adenocarcinoma": README.txt

Supplementary Material for Glycolysis-stratified coordination of fatty acid and glutamine metabolism in pancreatic ductal adenocarcinoma

This Zenodo repository contains all supplementary figures, tables, and source data referenced in the manuscript, including transcriptomic analyses (S1-S2), metabolite-inference analyses (S3-S5), a schematic of fatty-acid families and representative enzymes (S6), and proteomic validation (S7).

Contents:
- A Graphical Abstract 
S1 – Predictive Modeling of GLN Activity
S1 Fig. Elastic-Net out-of-fold predictions for GLN activity by glycolysis grouping. S1 Table. Cross-validated performance metrics (R² and RMSE) by grouping. S1 Data. Out-of-fold predicted vs. observed GLN activity by sample.

S2 – FA Family Definitions
S2 Table. Curated mapping of fatty-acid (FA) pathways/enzymes to functional families used in transcriptomic analyses.

S3 – Inferred Metabolite Activities
S3 Fig. Principal component analysis (PCA) of inferred metabolite activities across PDAC samples. S3 Table. Raw and z-scored inferred metabolite activity scores for 13 metabolites.

S4 – Metabolite State Comparisons
S4 Fig. Heatmap of mean z-scored metabolite activities across metabolite-defined metabolic phenotypes. S4 Table. One-way ANOVA and Kruskal–Wallis statistics comparing metabolite activities across phenotypes.

S5 – Mapping and Final State Labels
S5 Table. Metabolite–gene mapping with +1/−1 directionality assignments used to compute weighted metabolite activity scores. S5 Data. Final metabolite-defined metabolic state labels for all PDAC samples.

S6 – FA Chain-length Schematic and CPTAC Proteome Characteristics
S6 Fig. Schematic mapping representative enzymes and pathways across fatty-acid chain-length classes (MCFA, LCFA, VLCFA) and two example lipid families (glycerophospholipids, sphingolipids).
S6 Table. Descriptive statistics for the CPTAC PDAC tumor proteome (MD_abundance_tumor matrix), including number of tumors (n = 140), total proteins (11,662), per-protein missingness distribution, and the number of proteins retained after filtering to ≥70% observed values (7,993).

S7 – Proteomic Validation of FA–GLN Coordination (CPTAC PDAC Cohort)
S7 Fig. Proteomic validation of sphingolipid–glutamine coordination by glycolysis state in PDAC. Family-level associations between GLN and FA protein modules (glycerophospholipid, sphingolipid) are shown for low vs. high glycolysis tertiles.
S7 Table. Proteomic validation statistics for FA–GLN family-level correlations across glycolysis strata in the CPTAC PDAC cohort (n = 140). For each FA family and glycolysis tertile, the table reports sample size, Spearman correlation between GLN and FA protein-module scores, nominal p-values, and Benjamini–Hochberg FDR-adjusted q-values.

Contact: For questions regarding these supplementary files, please contact author.
