## Supplementary figures and images for "Glycolysis-stratified coordination of fatty acid and glutamine metabolism in pancreatic ductal adenocarcinoma"

### GA.png

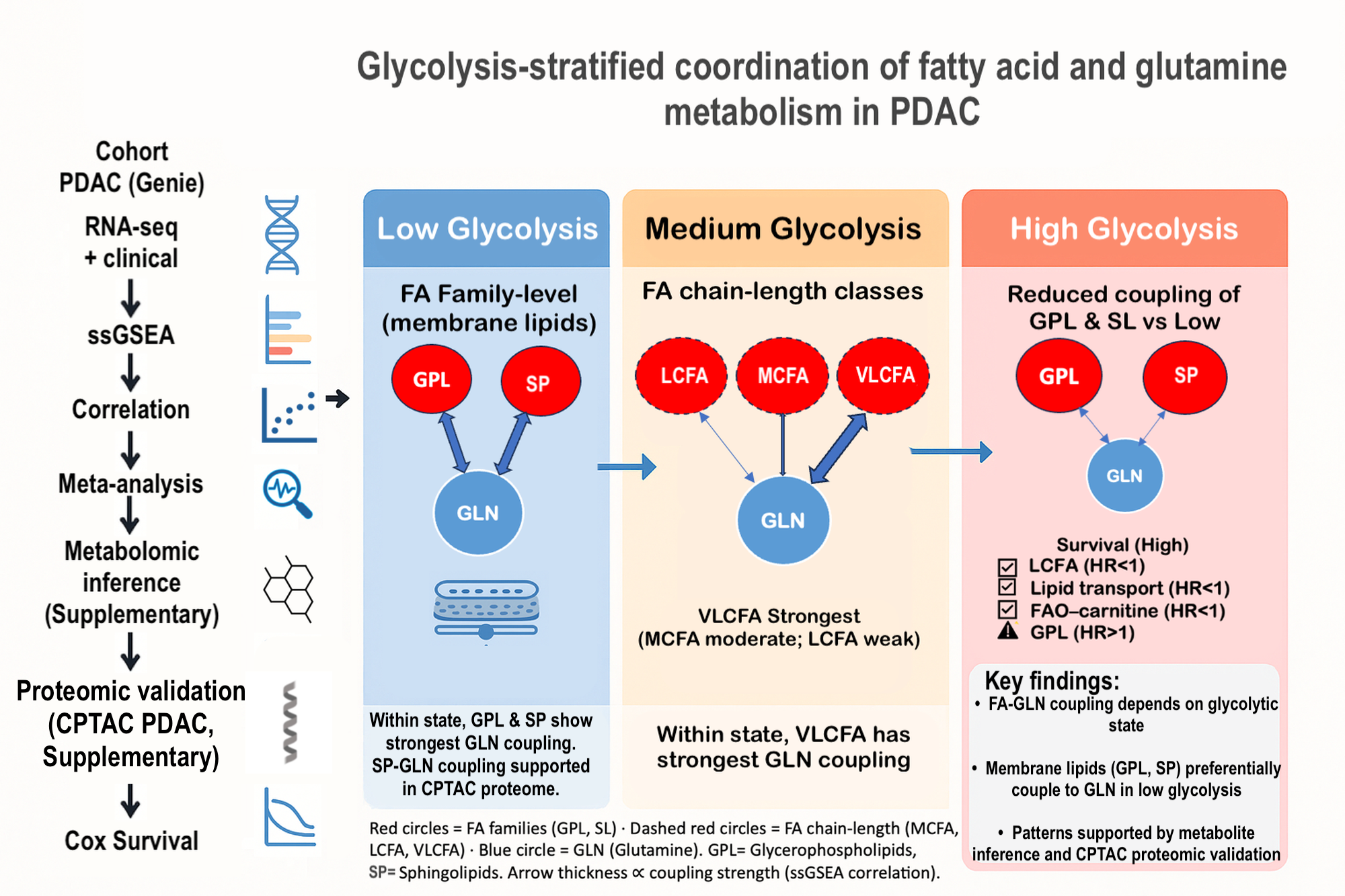

### S1_Fig_Enet_OOF.pdf

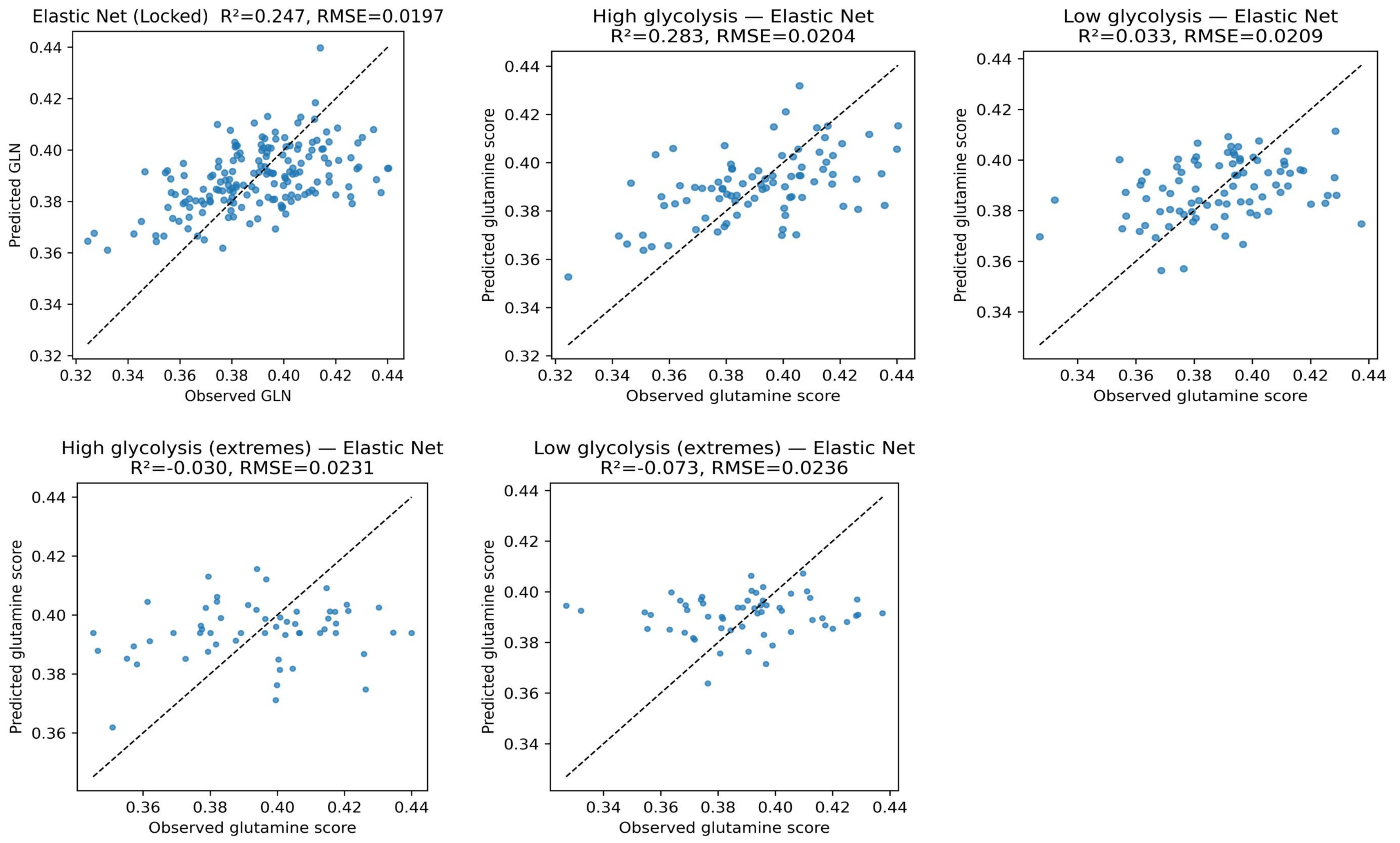

### S3_Fig_Metabolite_PCA.png

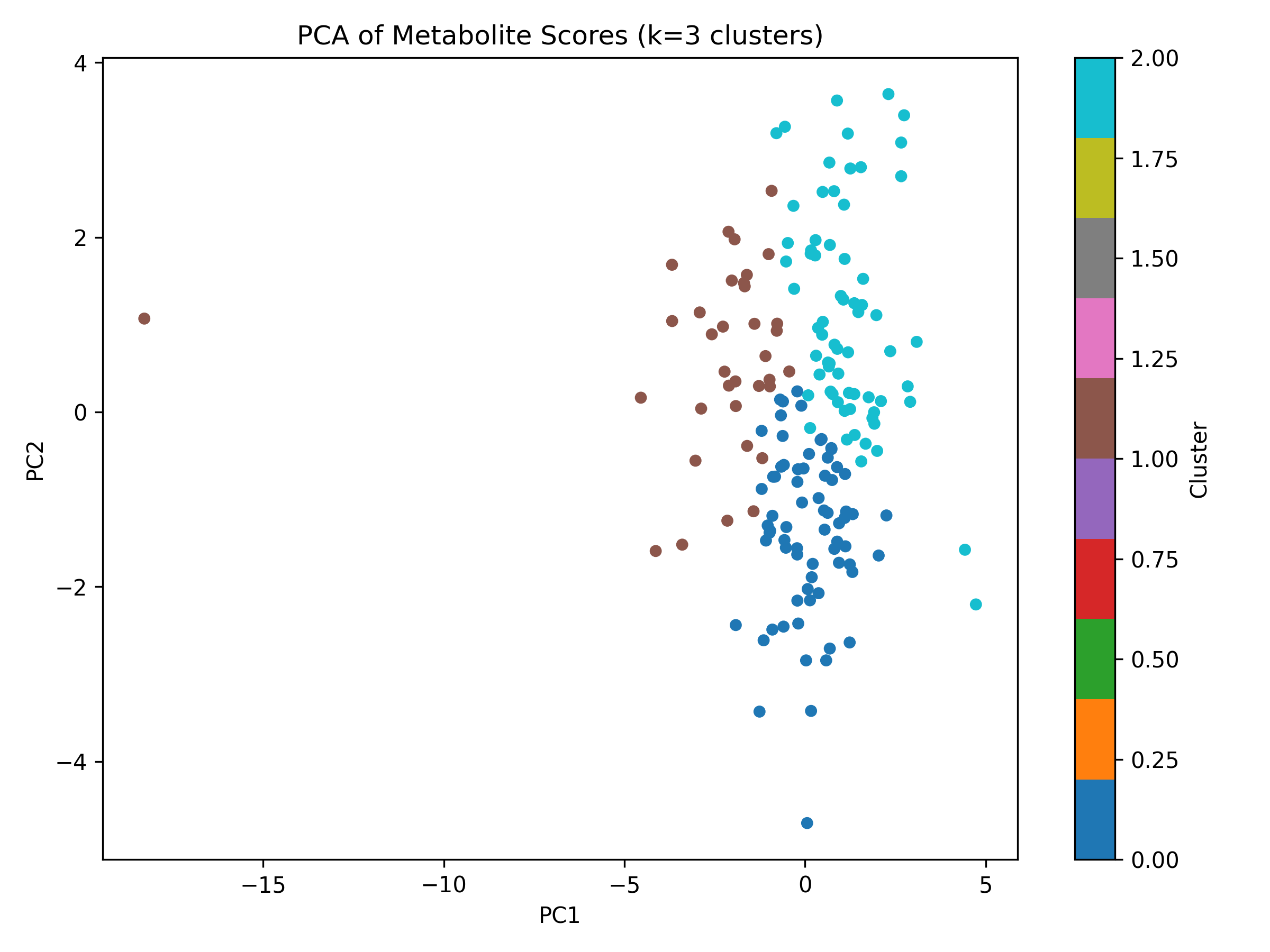

### S4_Fig_MetaboliteState_Heatmap.png

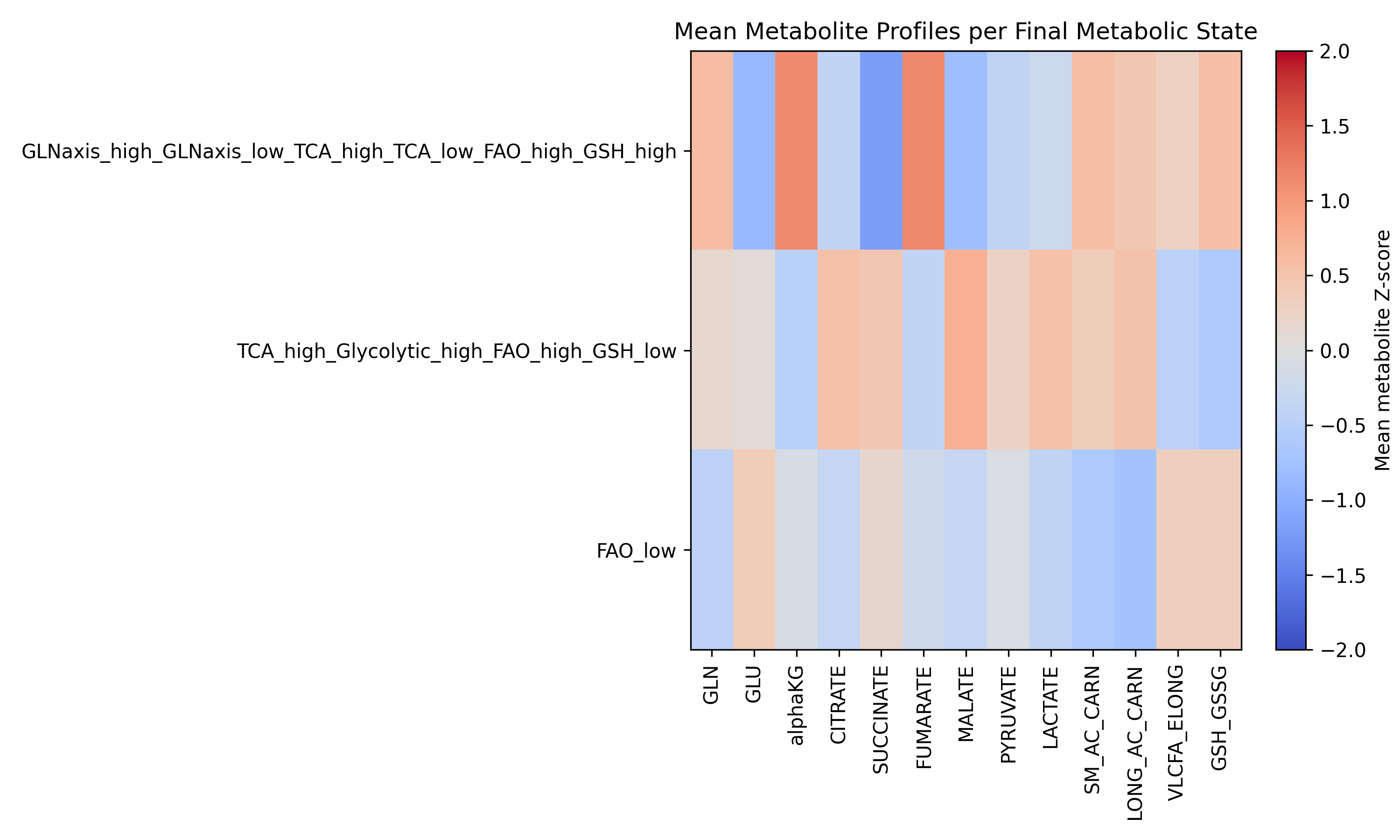

### S6_Fig_SchematicMapping_ChainLength_Family.png

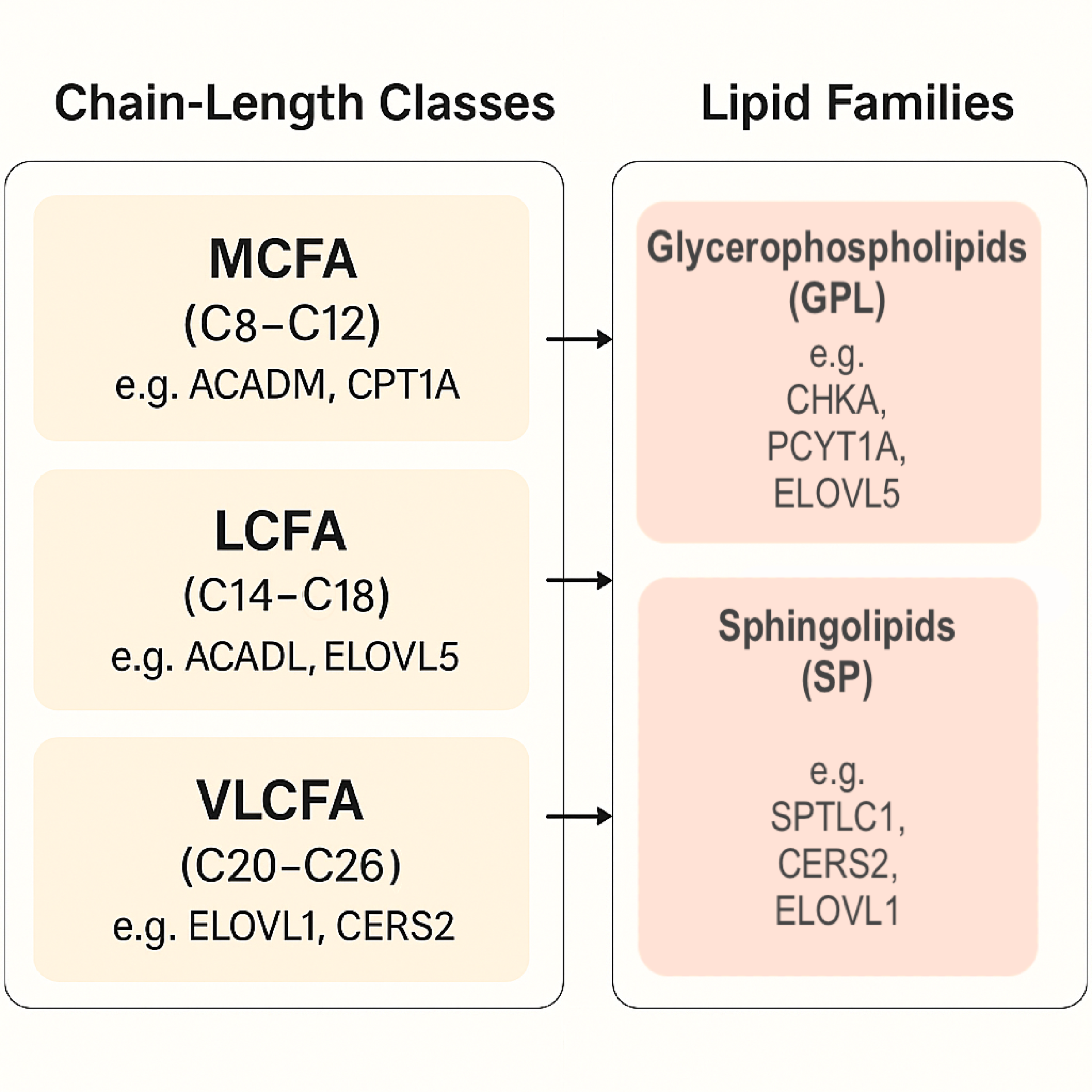

### S7_Fig_CPTAC_PDAC_FA_GLN_Family_Validation.png

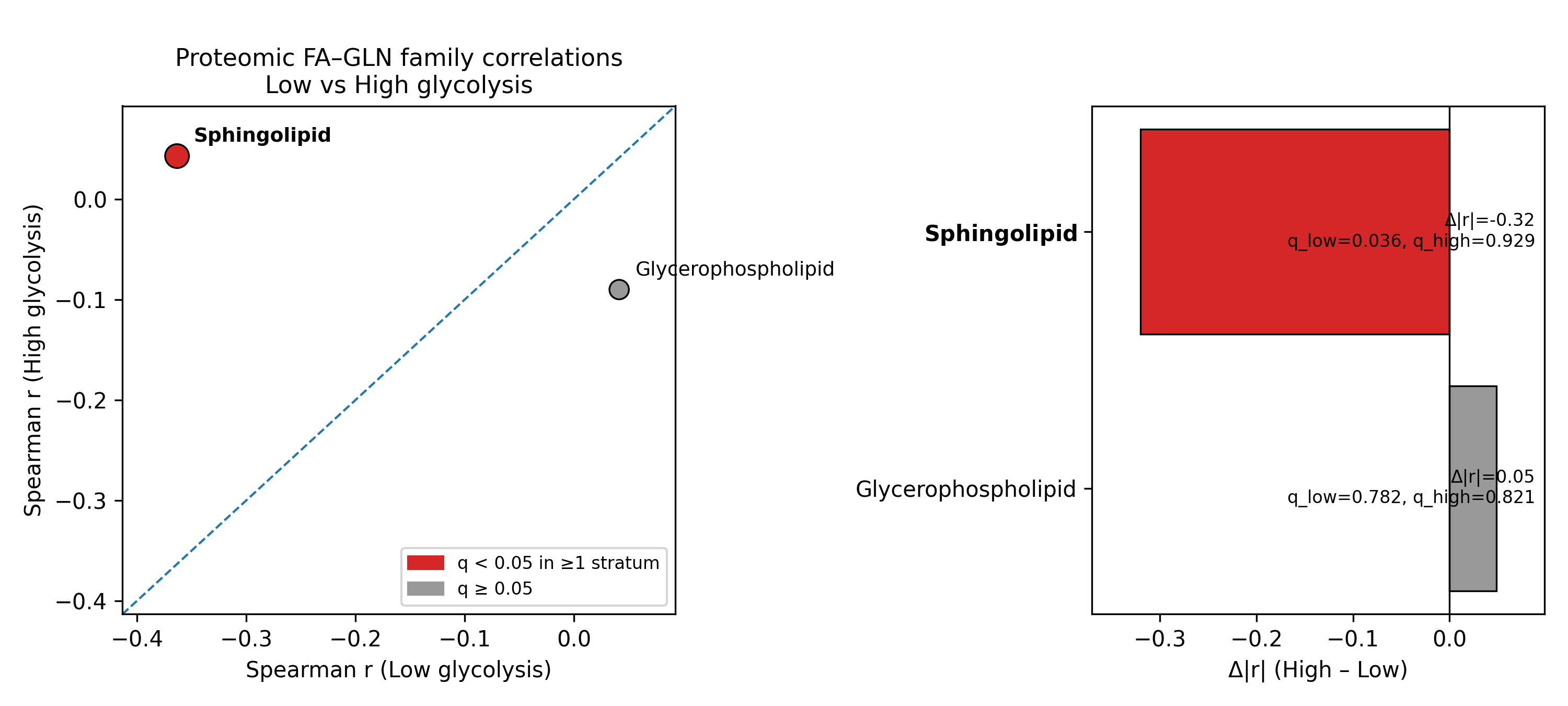
